# Using DNA metabarcoding to investigate diet and niche partitioning in the native European otter (*Lutra lutra*) and invasive American mink (*Neovison vison*)

**DOI:** 10.1101/2020.07.03.186346

**Authors:** Lynsey R. Harper, Hayley V. Watson, Robert Donnelly, Richard Hampshire, Carl D. Sayer, Thomas Breithaupt, Bernd Hänfling

**Author notes:** **Corresponding author:** Lynsey Harper, School of Natural Sciences and Psychology, Liverpool John Moores University, Liverpool, L3 3AF, UK.

## Abstract

In the UK, the native European otter (*Lutra lutra*) and invasive American mink (*Neovison vison*) have experienced concurrent declines and expansions. Currently, the otter is recovering from persecution and waterway pollution, whereas the mink is in decline due to population control and probable interspecific interaction with the otter. We explored the potential of DNA metabarcoding for investigating diet and niche partitioning between these mustelids. Otter spraints (*n* = 171) and mink scats (*n* = 19) collected from three sites (Malham Tarn, River Hull, and River Glaven) in northern and eastern England were screened for vertebrates using high-throughput sequencing. Otter diet mainly comprised aquatic fishes (81.0%) and amphibians (12.7%), whereas mink diet predominantly consisted of terrestrial birds (55.9%) and mammals (39.6%). The mink used a lower proportion (20%) of available prey (*n* = 40 taxa) than the otter, and low niche overlap (0.267) was observed between these mustelids. Prey taxon richness of mink scats was lower than otter spraints, and beta diversity of prey communities was driven by taxon turnover (i.e. the otter and mink consumed different prey taxa). Considering otter diet only, prey taxon richness was higher in spraints from the River Hull catchment, and beta diversity of prey communities was driven by taxon turnover (i.e. the otter consumed different prey taxa at each site). Studies using morphological faecal analysis may misidentify the predator as well as prey items. Faecal DNA metabarcoding can resolve these issues and provide more accurate and detailed dietary information. When upscaled across multiple habitat types, DNA metabarcoding should greatly improve future understanding of resource use and niche overlap between the otter and mink.

## Introduction

Dietary studies play a fundamental role in ecological research through revealing the feeding ecology of key species, the degree of resource overlap between species, and reconstructing complex trophic networks (Martínez-Gutiérrez et al. 2015). Morphological faecal analysis is a common method used to infer diet composition, especially in vertebrates. For example, morphological identification of prey item components from faeces has frequently been used to understand feeding ecology and resource overlap in mustelid predators, such as the European otter (*Lutra lutra*) and American mink (*Neovison vison*) (Jędrzejewska et al. 2001; Bonesi et al. 2004; Melero et al. 2008). However, morphological faecal analysis can be time-consuming, and accuracy hinges on possessing the necessary expertise to identify both the predator and its prey (Pompanon et al. 2012; Martínez-Gutiérrez et al. 2015; Traugott et al. 2020). Carnivore scats can be misidentified during field collection, with especially high error rates for sympatric species with morphologically similar scats and low density carnivores whose scats are sparse (Davison et al. 2002; Akrim et al. 2018). Prey detection from predator faeces may be influenced by differential digestion of soft-bodied and hard-bodied prey, and variable gut transition times for different prey components (e.g. hair, feather, teeth, bones, scales, shell) and prey types (e.g. fish, amphibian, bird, mammal) (Carss and Parkinson 1996; Krawczyk et al. 2016; Nielsen et al. 2018). Digestion can be influenced by species identity, life stage, and activity of predators as well as environmental variables (King et al. 2008; McInnes et al. 2016; Traugott et al. 2020). Smaller prey are less likely to be recovered from faeces, prey components may be fragmented or damaged beyond recognition, and prey components from related species can be morphologically similar. These issues individually or combined can prevent species-level identification for various taxa, especially fishes (e.g. closely related cyprinids) and birds (Britton et al. 2006, 2017; Shehzad et al. 2012a; Krawczyk et al. 2016; Berry et al. 2017; Smiroldo et al. 2019; Traugott et al. 2020).

Molecular tools offer a rapid, non-invasive, cost-efficient alternative to morphological faecal analysis for identification of predator and prey. Single or multiple prey species within a taxonomic group can be targeted using species- or group-specific DNA barcodes, or prey species across multiple taxonomic groups can be assessed in parallel using generic DNA metabarcodes with high-throughput sequencing, i.e. DNA metabarcoding (Pompanon et al. 2012; McInnes et al. 2016; Traugott et al. 2020). DNA metabarcoding cannot provide information on cannibalism, or size, life stage, and vital status of prey taxa, and is not immune to retention of prey taxa due to differential digestion and gut transition times. Nonetheless, it perpetuates non-invasiveness and has greater sensitivity toward rare, soft, liquid or highly degraded prey items, e.g. jellyfish in faeces of marine predators (Shehzad et al. 2012b; McInnes et al. 2017; Nielsen et al. 2018; Traugott et al. 2020). As such, DNA metabarcoding provides species resolution data at greater spatiotemporal scales for the vast majority of prey items, regardless of prey size, type, and integrity or lack of hard components (Oehm et al. 2011; Pompanon et al. 2012; McInnes et al. 2017; Forin-Wiart et al. 2018; Traugott et al. 2020). Since its inception, DNA metabarcoding has been employed to assess the diet of various mammalian predators (Shehzad et al. 2012a, 2012b; De Barba et al. 2014; Berry et al. 2017; Gosselin et al. 2017; Forin-Wiart et al. 2018; Robeson et al. 2018; Schwarz et al. 2018), and recent small-scale studies have shown its potential for European otter (hereafter otter) diet analysis (Buglione et al. 2020; Martínez-Abraín et al. 2020).

Dietary niche characterisation of the otter is important as this is a keystone species and an apex predator of freshwater ecosystems in Europe. In the UK, the otter was common and widespread until the 18th century, after which the population declined sharply due to persecution, bioaccumulation of polychlorinated biphenyls (PCBs), and organochlorine pesticide poisoning, resulting in local extinctions over large tracts of its former range (Britton et al. 2006; McDonald et al. 2007; Harrington et al. 2009; Reid et al. 2013; Smiroldo et al. 2019). However, legal protection, pesticide bans, water quality and habitat improvement, and targeted otter releases since the 1980s allowed the species to recover (Bonesi and Macdonald, 2004a; Britton et al. 2006; McDonald et al. 2007; Alderton et al. 2015; Martínez-Abraín et al. 2020). Conversely, the American mink (hereafter mink) was introduced from America to Europe for fur farming in the 1920s, and became established in the wild and invasive across Europe following fur farm escapees and intentional releases (Bonesi and Macdonald 2004b; Reynolds et al. 2004; Bonesi and Palazon 2007; Harrington et al. 2009). In the UK, rapid countrywide spread of the mink has been documented since the 1950s (Bonesi and Macdonald 2004b; Reynolds et al. 2004). This mustelid has had acutely devastating effects on native UK biodiversity, including the European water vole (*Arvicola amphibius*) and ground-nesting seabirds (Bonesi and Macdonald 2004a, 2004b; Reynolds et al. 2004), due to direct predation. The species has also proven economically damaging, with poultry runs, gamebird rearing, and fisheries all negatively affected by mink activity (Bonesi and Palazon 2007).

Initially, there was misplaced belief that the mink had contributed to the decline of the otter through competition due to simultaneous changes in distribution and abundance of these two mustelids (McDonald et al. 2007). However, studies on interspecific aggression and intraguild predation have shown that the otter is more likely to be the victor in encounters between these mustelids due to its larger body size, heavier weight, and better swimming/diving skills (Bonesi and Macdonald 2004a; Bonesi et al. 2004; Melero et al. 2008; Harrington et al. 2009). Furthermore, declines in mink site occupancy and density have been linked to otter recovery at fine and broad spatiotemporal scales (Bonesi and Macdonald 2004a; McDonald et al. 2007). The otter has been classed as a specialist or generalist predator, whereas the mink is typically considered to be an opportunist (Jędrzejewska et al. 2001; Bonesi and Macdonald 2004b; Melero et al. 2008; Almeida et al. 2012, 2013; Reid et al. 2013). Evidence indicates that the otter outcompetes the mink for aquatic prey, resulting in the mink seeking out terrestrial prey and undergoing a feeding niche shift where these mustelids are sympatric (Jędrzejewska et al. 2001; Bonesi et al. 2004; Melero et al. 2008; Harrington et al. 2009). Indeed, niche overlap between the otter and mink has been found to be lower in winter than spring, possibly due to restricted resources (Jędrzejewska et al. 2001; Bonesi et al. 2004). Additionally, both species have been found to consume different prey in response to water body type and size (Jędrzejewska et al. 2001). Coexistence of these two species is highly dependent on riparian habitat features and terrestrial prey availability, but dietary and spatial segregation of the otter and mink can eventually occur (Bonesi and Macdonald 2004a, 2004b; Harrington et al. 2009). It is unknown whether this niche partitioning may exacerbate mink predation of native and threatened UK biodiversity.

We assessed the potential of DNA metabarcoding for investigating dietary profiles of the native otter and invasive mink, and resource overlap between these mustelids. Otter spraints and mink scats were collected at three study sites across northern and eastern England: Malham Tarn, a calcareous upland lake in North Yorkshire; River Glaven, a lowland chalk stream in North Norfolk; and the River Hull, a chalk stream in East Yorkshire. DNA extracted from faecal matter was analysed for all vertebrate species using high-throughput sequencing. We hypothesised low resource overlap between the otter and mink. The otter was expected to predate a broad range of aquatic and semi-aquatic prey (i.e. fish, amphibians, waterfowl) whereas the mink was anticipated to specialise on semi-aquatic and terrestrial species (i.e. birds, mammals) as documented by studies that used morphological faecal analysis.

## Methods

### Study sites and sample collection

Mammal faeces were collected from 2015 to 2018 in northern and eastern England: River Hull catchment, East Yorkshire (sites along the river and ponds in close proximity to the river); Malham Tarn (lake) and Gordale Beck (stream close to Malham Tarn), West Yorkshire; and River Glaven catchment, Norfolk (sites along the river and ponds in close proximity) (Fig. S1). Sample information, including collection date, coordinates, and site, is provided in Table S1. Faeces were ostensibly identified as otter spraints (*n* = 206), mink scats (*n* = 9), and red fox (*Vulpes vulpes*) scat (*n* = 1). The red fox scat was collected despite being a non-focal mammal predator due to potential predator misidentification using faecal characteristics. Faeces were collected using zip-lock bags or 50 mL falcon tubes (SARSTEDT, Germany, UK) and frozen at −20 °C until DNA extraction. For each site, a basic inventory of fishes was created from available survey data to permit a broad comparison between prey detected in otter spraints by DNA metabarcoding and available prey species (Supplementary Material: Appendix 1). Fish survey (seine netting, electrofishing) data from 2000 to 2019 were extracted from the publicly available Environment Agency database (data.gov.uk) for the River Hull and River Glaven catchments. For the River Glaven, additional data were available from the surveys of Harwood et al. (2019) and Sayer et al. (in press). Fish community data for Malham Tarn were obtained through environmental DNA (eDNA) metabarcoding verified by fishery owners (Hänfling et al. 2020) and from fish surveys detailed in Eldridge (2016).

### DNA extraction

DNA was extracted from faeces using the DNeasy PowerSoil Kit (Qiagen^®^, Hilden, Germany) or the Mu-DNA soil protocol with a tissue protocol wash stage (Sellers et al. 2018). Using a bleach and ultraviolet (UV) sterilised spatula and weigh boat (Merck, Darmstadt, Germany) for each sample, ≈0.25 g of faecal matter was measured out and added directly to pre-labelled PowerBead tubes for DNeasy PowerSoil extraction or 5 mL tubes (Axygen™, Fisher Scientific, UK) containing 0.5 g of 1-1.4 mm diameter sterile garnet beads (Key Abrasives Ltd., UK) for Mu-DNA extraction. Either 60 μL of Solution C1 (DNeasy PowerSoil) or 550 μL Lysis Solution and 200 μL Soil Lysis Additive (Mu-DNA) was added to each tube. Tubes were placed in a Qiagen^®^ TissueLyser (30 frequencies/min) for 10 min to homogenise the samples. Remaining steps were performed according to the DNeasy PowerSoil or Mu-DNA protocol. Eluted DNA (100 μL) concentration was quantified on a Qubit™ 3.0 fluorometer using a Qubit™ dsDNA HS Assay Kit (Invitrogen, UK). DNA extracts were stored at −20 °C prior to PCR.

### DNA metabarcoding

Samples were processed for DNA metabarcoding in two libraries. One library contained the samples from the River Hull catchment, collected between 2015 and 2017, while the other library contained samples from the River Hull, River Glaven and Malham Tarn collected in 2018. DNA metabarcoding followed the procedures established by Harper et al. (2019a) which are described in Supplementary Material: Appendix 2. Briefly, double-indexed libraries were constructed with a two-step PCR protocol which first used published primers 12S-V5-F (5’-ACTGGGATTAGATACCCC-3’) and 12S-V5-R (5’-TAGAACAGGCTCCTCTAG-3’) with modifications (i.e. indexes, heterogeneity spacers, sequencing primers, and pre-adapters) to amplify a region of the 12S ribosomal RNA (rRNA) mitochondrial gene (Riaz et al. 2011). These primers have been validated *in silico*, *in vitro*, and *in situ* for UK vertebrates (Hänfling et al. 2016; Harper et al. 2019a, 2019b). Exotic cichlid (*Maylandia zebra*) DNA (0.05 ng/μL) was the PCR positive control, and sterile molecular grade water (Fisher Scientific UK Ltd, Loughborough, UK) was the PCR negative control. Three PCR replicates were performed for each DNA sample and pooled prior to normalisation. Normalised sub-libraries were created by pooling PCR products according to band strength and PCR plate, and purified with Mag-BIND^®^ RxnPure Plus magnetic beads (Omega Bio-tek Inc, GA, USA) following a double size selection protocol (Bronner et al. 2009). PCR in duplicate bound pre-adapters, indexes, and Illumina adapters to the purified sub-libraries, and PCR replicates were pooled for magnetic bead purification. Sub-libraries were quantified on a Qubit™ 3.0 fluorometer using a Qubit™ dsDNA HS Assay Kit, and pooled proportional to sample size and concentration for magnetic bead purification. An Agilent 2200 TapeStation and High Sensitivity D1000 ScreenTape (Agilent Technologies, CA, USA) were used to verify fragment size (330 bp) of the final libraries and absence of secondary product. The libraries were quantified using real-time quantitative PCR with the NEBNext^®^ Library Quant Kit for Illumina^®^ (New England Biolabs^®^ Inc., MA, USA) on a StepOnePlus™ Real-Time PCR system (Life Technologies, CA, USA) and diluted to 4 nM. Each library (one containing 125 faecal samples and eight PCR controls, and one containing 140 faecal samples, 12 PCR controls, and 12 external samples) was sequenced at 12 pM with 10% PhiX Control v3 on an Illumina MiSeq^®^ using a MiSeq Reagent Kit v3 (600-cycle) (Illumina Inc., CA, USA). Raw sequence reads were demultiplexed with a custom Python script. Sequences underwent quality trimming, merging, chimera removal, clustering, and taxonomic assignment against our custom reference database for UK vertebrates (Harper et al. 2019b) using metaBEAT v0.97.11 (https://github.com/HullUni-bioinformatics/metaBEAT). Taxonomic assignment used a lowest common ancestor approach based on the top 10% BLAST matches for any query that matched a reference sequence across >80% of its length at a minimum identity of 98%. Unassigned sequences were compared against the NCBI nucleotide (nt) database at 98% minimum identity using the same lowest common ancestor approach. The bioinformatic analysis has been deposited in a dedicated GitHub repository, which has been permanently archived for reproducibility (https://doi.org/10.5281/zenodo.4252552)

### Data analysis

Analyses were performed in the statistical programming environment R v.3.6.3 (R Core Team, 2020) unless otherwise stated. Data and R scripts have been deposited in the GitHub repository. Dataset refinement is summarised here and fully described in Supplementary Material: Appendix 2. BLAST results from different databases were combined and spurious assignments were removed. Where applicable, orders, families and genera containing a single UK species were reassigned to that species, species were reassigned to domestic subspecies, and misassignments were corrected. The read counts for metaBEAT and manual assignments were merged prior to application of a sequence threshold (i.e. maximum sequence frequency of cichlid DNA in faecal samples) to mitigate against contamination and false positives in the dataset (Figs S2, S3). After applying the false positive threshold (1.123%), taxonomic assignments above species-level were removed with exceptions (Supplementary Material: Appendix 2). Human (*Homo sapiens*) and domestic animals (cow [*Bos taurus*], dog [*Canis lupus familiaris*], pig [*Sus scrofa domesticus*]) were regarded as environmental contaminants and also removed for the purposes of downstream analyses.

Using Microsoft Excel, each faecal sample was assigned to a mammal predator based on the proportional read counts for each predator species (otter, mink, red fox and European polecat [*Mustela putorius*]) detected (Supplementary Material: Appendix 3). In cases where DNA from multiple predators was present, the sample was assigned to the predator species which possessed more than 90% of the total predator read counts. If no predator species possessed more than 90% of the total predator read counts in a sample or a sample contained less than 100 reads for all predators, the sample was removed from the dataset. After predator assignment, the total percentage of prey (by vertebrate group) sequences relative to predator sequences was evaluated across all samples belonging to each predator in R (otter and mink) or Microsoft Excel (red fox and European polecat; Appendix 4). Using R, all predator reads, and samples belonging to red fox (hereafter fox) and European polecat (hereafter polecat), were then removed for downstream analyses.

In R, the data for otter and mink samples were summarised as the total percentage of prey sequences for each vertebrate group, proportional read counts for each prey taxon in each sample, and the percentage frequency of occurrence (i.e. the percentage of faecal samples that a prey taxon was detected in). The read count data were converted to presence/absence using the DECOSTAND function in the package vegan v2.5-6 (Oksanen et al. 2018). We used the package bipartite v2.15 (Dormann et al. 2009) to construct a semi-quantitative trophic network for each predator and their prey. Network-level metrics were obtained using the NETWORKLEVEL function, and species-level metrics for each predator obtained using the SPECIESLEVEL function. Taxon richness (alpha diversity) was obtained using the SPECNUMBER function in the package vegan v2.5-6. Given that the data were not normally distributed (Shapiro-Wilk normality test: W = 0.921, *P* < 0.001) and the number of samples between predators and sampling locations was unbalanced, Kruskal-Wallis tests followed by Dunn’s tests, from the packages stats v3.6.3 and FSA v0.8.30 (Ogle et al., 2020) respectively, were used to compare alpha diversity of prey communities between otter and mink faecal samples, and between otter spraints from different sites. Data for the mink and each freshwater habitat were too sparse for examination of geographic variation in mink diet, and differences in otter and mink diet with regard to habitat (Fig. S4). We used the package iNEXT v2.0.20 (Hsieh et al. 2016) to perform rarefaction and extrapolation curves to ensure that differences in prey taxon richness were not driven by imbalances in sample size for predators and sampling locations. The INEXT function was run using incidence frequencies for prey taxa with 300 samples, 60 knots, 1000 bootstraps, and 95% confidence intervals. The ESTIMATED function was used to perform both sample size-based and coverage-based comparisons between predators and sampling sites (otter only) with 95% confidence intervals and 95% sample coverage (coverage-based comparison only).

Before partitioning beta diversity, we compared prey community dissimilarity inferred from occurrence (i.e. presence/absence) and relative read abundance (RRA; i.e. proportional read counts) data. Using the package vegan v2.5-6, the read count data were converted to presence/absence and proportional read count matrices using the DECOSTAND function. Jaccard and Bray-Curtis dissimilarity indices were computed for the presence/absence and proportional read counts matrices respectively using the VEGDIST function, and beta diversity was visualised using Non-metric Multidimensional Scaling (NMDS) with the METAMDS function. Two outlier samples containing one or two taxa were removed to improve visualisation of variation in otter and mink diet (LIB02-TL01 [mink] and LIB02-TL07 [otter]) and site variation in otter diet (LIB02-TL07 and LIB04-TL57), but patterns produced by occurrence and RRA data were comparable (Fig. S5). Given that our stringent false positive sequence threshold should have removed any minor prey items, secondary predation, and contaminants, we used occurrence data with Jaccard dissimilarity for beta diversity partitioning to mitigate potential taxon recovery bias (Deagle et al. 2018).

We employed the package betapart v1.5.1 (Baselga and Orme 2012) to estimate total beta diversity, partitioned by turnover (i.e. community dissimilarity due to taxon replacement) and nestedness-resultant (i.e. community dissimilarity due to taxon subsets), with the BETA.MULTI (multiple-site dissimilarities) and BETA.PAIR (pairwise dissimilarity matrices) functions. For each component of beta diversity, we compared community heterogeneity in faecal samples grouped by predator (otter or mink) or site of otter spraint collection (Malham Tarn, River Glaven, River Hull) by calculating homogeneity of multivariate dispersions (MVDISP) using the BETADISPER function from the package vegan v2.5-6. Variation in MVDISP between otter and mink samples or between otter spraints from different sites was statistically tested using ANOVA. Differences in prey communities for each component of beta diversity were visualised using NMDS with the METAMDS function, and tested statistically using permutational multivariate analysis of variance (PERMANOVA) with the function ADONIS in the package vegan v2.5-6. Pre-defined cut-off values were used for effect size, where PERMANOVA results were interpreted as moderate and strong effects if R^2^ > 0.09 and R^2^ > 0.25 respectively. These values are broadly equivalent to correlation coefficients of *r* = 0.3 and 0.5 which represent moderate and strong effects accordingly (Nakagawa & Cuthill, 2007). All figures were produced using the package ggplot2 v3.3.1 (Wickham, 2009), except Fig. 4 which was produced in Microsoft PowerPoint. Legends for Figs 1, 2, and 3 were adjusted using Inkscape (http://www.inkscape.org/).

**Figure 1.**
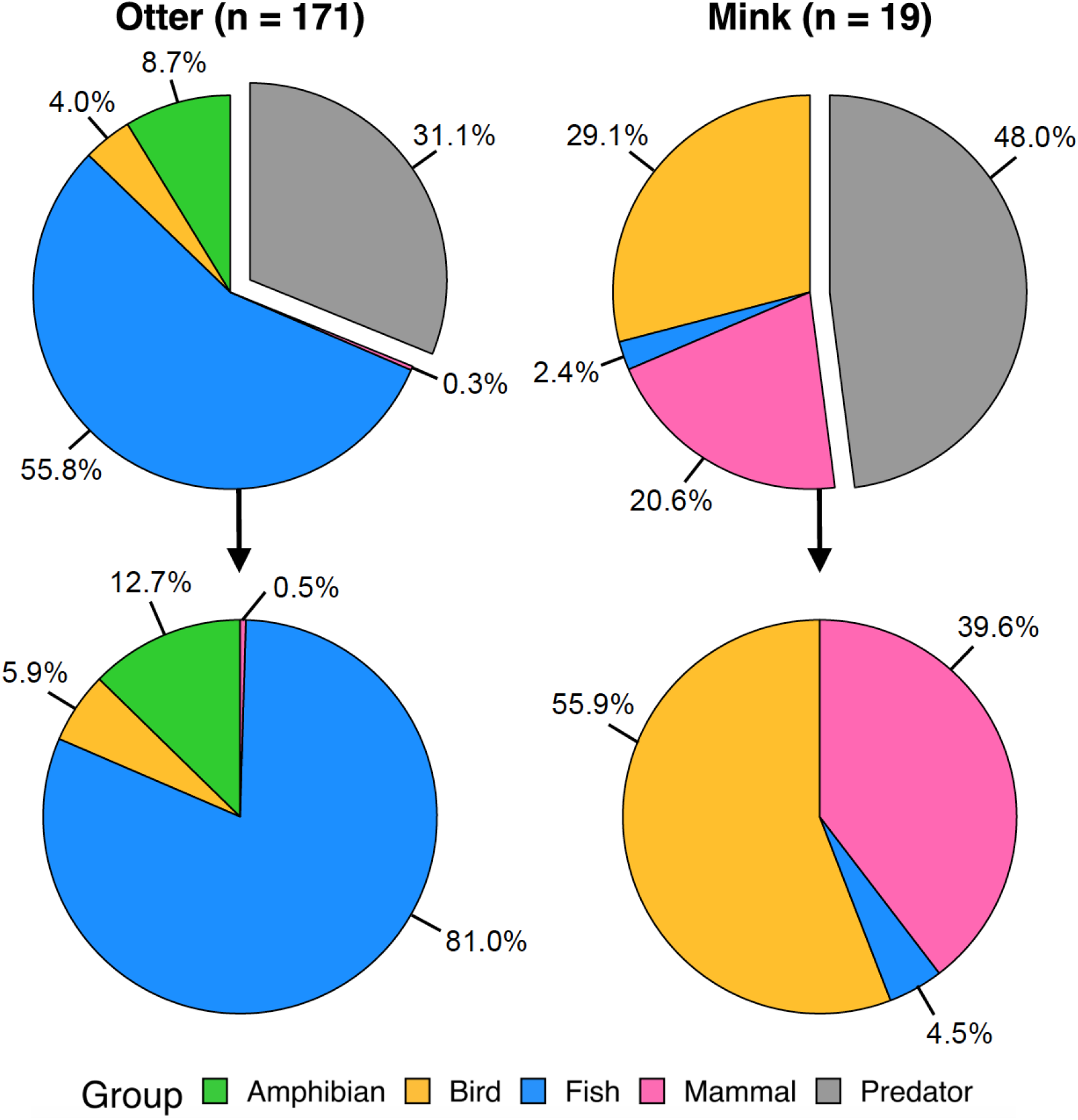
Pie charts showing the proportion of total reads retained in the refined dataset that belonged to the otter and mink with respect to their vertebrate prey, and the proportion of prey reads that belonged to different vertebrate groups.

**Figure 2.**
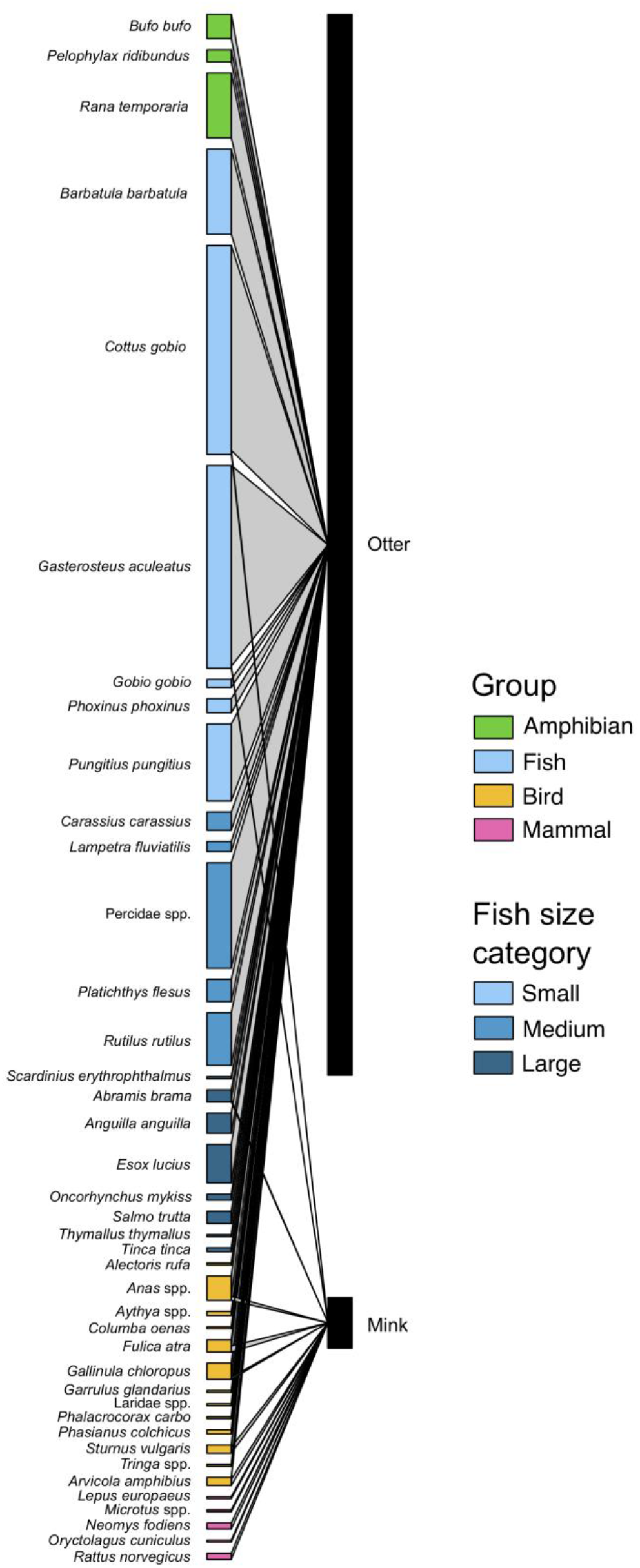
A bipartite trophic network showing the prey of the otter and mink. The black blocks on the right column represent the predators and the coloured blocks in the left column represent the prey taxa. Detected predation events are indicated by lines that connect a predator with a prey taxon, and the number of events is proportional to the thickness of the line. Prey taxa are coloured according to vertebrate group, and different shades of blue indicate fish size category.

**Figure 3.**
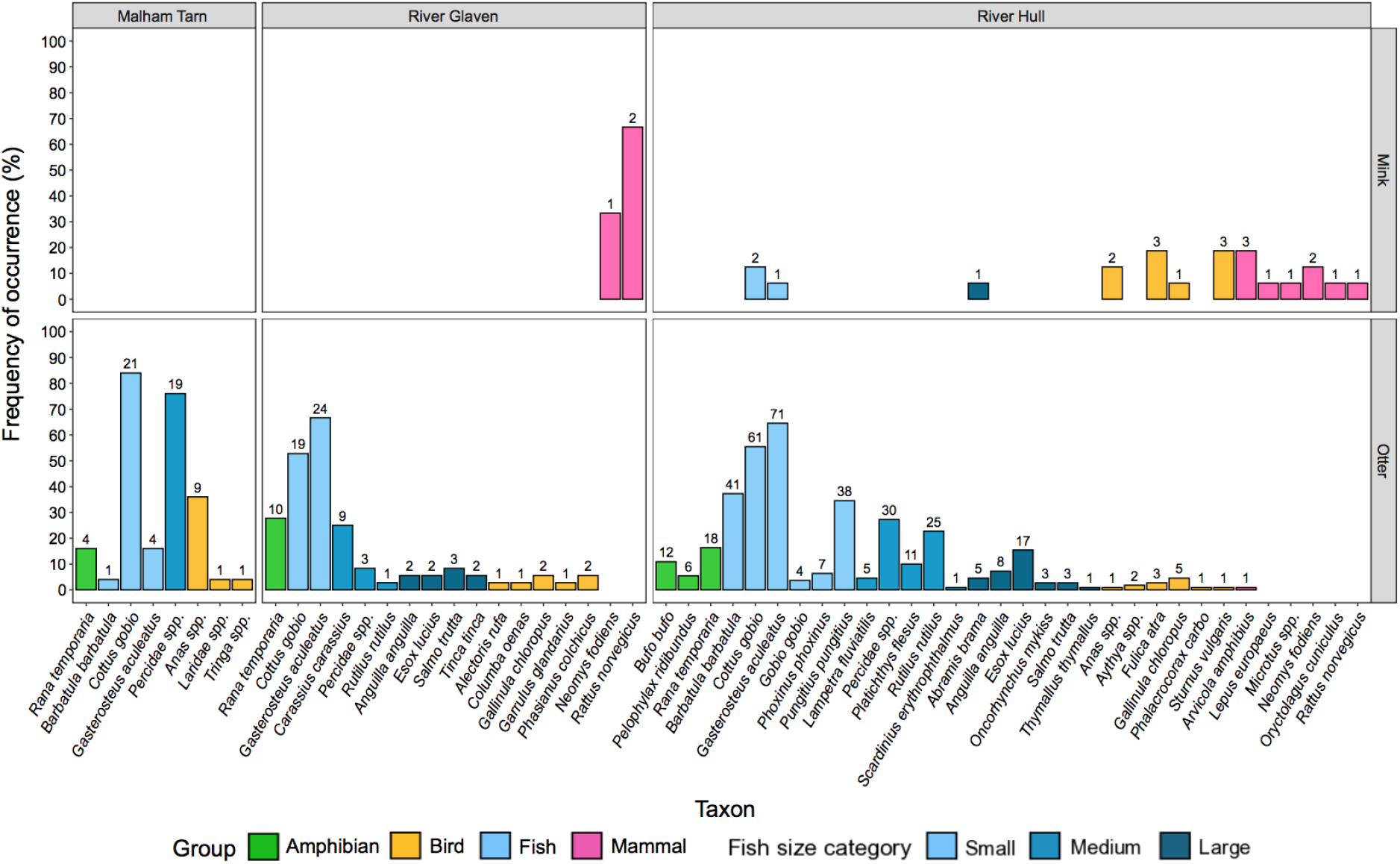
Barplot showing the occurrence percentage of prey taxa in mink and otter samples collected from different sites. Bars are coloured according to vertebrate group, and different shades of blue indicate fish size category. Numbers above bars represent the number of samples where prey taxa were detected.

**Figure 4.**
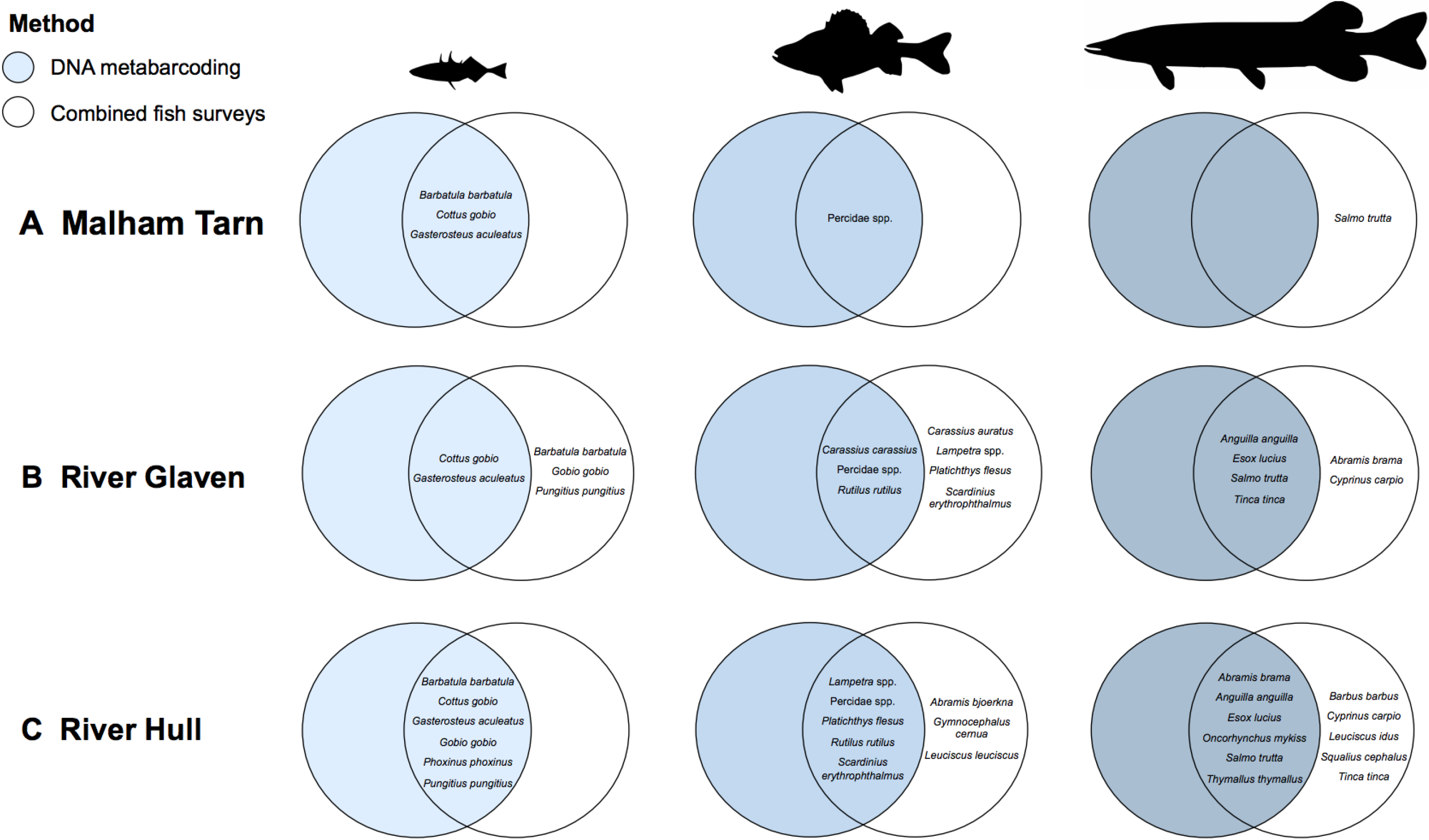
Venn diagrams showing fish species belonging to different size categories that were detected by DNA metabarcoding of otter spraints (blue circles) or fish surveys (white circles) at **A** Malham Tarn, **B** River Glaven, and **C** River Hull.

## Results

### Data filtering

The libraries generated a total of 22,286,976 and 40,074,340 raw sequence reads respectively, which were reduced to 9,487,780 and 14,362,257 reads by trimming, merging, and length filter application. After removal of chimeras and redundancy via clustering, 9,340,695 and 14,153,929 reads remained (average read count of 72,408 and 86,304 per sample including controls), of which 9,244,260 (98.97%) and 13,909,558 (98.27%) were assigned a taxonomic rank. Contamination from different sources was observed in the PCR controls (Fig. S2) as well as cichlid DNA in the faecal samples. No cichlid DNA remained in the faecal samples after application of the false positive sequence threshold, and taxonomic assignments were narrowed (Fig. S3). Before threshold application, we detected 127 taxa from 216 faecal samples, including six amphibian taxa, 43 fish taxa, 36 bird taxa, and 41 mammal taxa. However, 61 taxa (including two amphibian taxa, 20 fish taxa, 17 bird taxa, and 21 mammal taxa) were consistently detected below our threshold and were therefore removed from the dataset. The final dataset after threshold application and refinement of taxonomic assignments contained 46 taxa (38 assigned to species-level): three amphibians, 19 fishes, 13 birds, and 11 mammals.

### Predator assignment

Thirteen faecal samples contained less than 100 reads for any mammal predator and were removed from the dataset. In most of the remaining samples, DNA from a single predator comprised 100% of the total predator read counts (otter: *n* = 169; mink: *n* = 17; fox: *n* = 5; polecat: *n* = 1). Four samples with read counts for multiple predator species were assigned to a predator species based on a majority rule, i.e. the predator species possessed >90% of the total predator read counts (otter: *n* = 2; mink: *n* = 2). Seven samples were discarded because a confident predator assignment could not be made, i.e. no predator possessed >90% of the total predator read counts. Consequently, the refined dataset contained 171 otter, 19 mink, 5 fox, and 1 polecat faecal sample(s). For 90.82% of samples that were retained (*n* = 196), predator assignment was in agreement with visual identification of faeces. Predator assignment in 18 samples (9.18%) changed based on DNA metabarcoding. Fox and polecat diet is reported in Supplementary Material: Appendix 5.

### Otter and mink diet

Otter DNA and mink DNA encompassed 31.1% and 48.0% respectively of reads obtained from faecal samples belonging to these mustelids (Fig. 1). Using the prey reads, otter diet was mainly composed of fishes (81.0%) and amphibians (12.7%), whereas mink diet predominantly consisted of birds (55.9%) and mammals (39.6%) (Fig. 1).

The bipartite trophic network for the otter and mink contained 40 prey species (Fig. 2), of which eight were predated by both mustelids: bream (*Abramis brama*), European bullhead (*Cottus gobio*), three-spined stickleback (*Gasterosteus aculeatus*), ducks (*Anas* spp.), Eurasian coot (*Fulica atra*), common moorhen (*Gallinula chloropus*), starling *(Sturnus vulgaris*), and water vole (*Arvicola amphibius*) (Figs 2, 3). Notably, occurrence of mink predation on bream (5.26%), duck species (10.53%), Eurasian coot (15.79%), common moorhen (5.26%), starling (15.79%), and water vole (15.79%) was more frequent than occurrence of otter predation on these species (2.92%, 5.85%, 1.75%, 4.09%, 0.59%, and 0.59% respectively) (Fig. 3). Network-level metrics indicated some degree of specialisation (specialisation index *H2′* = 0.628), with few prey interactions for each predator (generality = 14.333) and a low proportion of possible interactions realised in the network (weighted connectance = 0.184), leading to few shared prey species between otter and mink (niche overlap = 0.267).

Species-level metrics for each predator provide further evidence for predator specialisation within the network. Both predators’ diets were relatively specialised (Paired Differences Index: otter = 0.893, mink = 0.812), but mink diet showed greater divergence from random selections of prey species (d’: otter = 0.526, mink = 0.671), with a lower proportion of available resources utilised (proportional similarity: otter = 0.962, mink = 0.209; unused resource range: otter = 0.128, mink = 0.692). However, resources within each predators’ diet were used relatively evenly, with neither species relying predominantly on a few key resources (species specificity index: otter = 0.287, mink = 0.267). Shannon diversity of predator-prey interactions was higher for the otter than the mink (partner diversity: otter = 2.672, mink = 2.449), suggesting that mink diet was less diverse. Only 13 prey species were detected in mink scats compared with 35 prey species in otter spraints (Figs 2, 3).

Prey species unique to the mink were brown hare (*Lepus europaeus*), *Microtus* spp., water shrew (*Neomys fodiens*), European rabbit, and brown rat (*Rattus norvegicu*s), but many fishes and amphibians were unique to the otter (Figs 2, 3, S6). Otter predation events largely involved common frog (*Rana temporaria*) and small, abundant fishes, such as European bullhead, stone loach (*Barbatula barbatula*), three-spined stickleback, and ninespine stickleback (*Pungitius pungitius*), with predation on medium (e.g. crucian carp [*Carassius carassius*], roach [*Rutilus rutilus*], Percidae spp.) and large (e.g. European eel [*Anguilla anguilla*], Northern pike [*Esox lucius*]) fishes occurring less frequently (Figs 3, S6). At each site, all fishes detected by DNA metabarcoding of otter spraints had also been recorded during recent surveys (conducted between 2000 and 2019) that used conventional fish monitoring tools or eDNA metabarcoding (Fig. 4). However, some fishes detected during previous surveys of the River Glaven (*n* = 9), River Hull (*n* = 8), and Malham Tarn (*n* = 1) were not found with faecal DNA metabarcoding (Fig. 4).

Two otter and two mink samples did not contain any prey taxa and were removed from the dataset for alpha and beta diversity analyses. Predator influenced alpha diversity of faecal samples (χ^2^_1_ = 22.786, *p* < 0.001), with taxon richness of mink scats significantly lower (*Z* = −4.773, *p* < 0.001) than that of otter spraints (Fig. 5a). Rarefaction and extrapolation curves indicated that lower prey taxon richness of mink scats was not due to disparities in sample size between predators. Prey taxon richness began to plateau at 21 taxa with 95 or more mink scats. In contrast, prey taxon richness did not plateau even with 300 otter spraints, at which 42 taxa would be detected (Fig. 5bi). Over 1100 otter spraints would be required for prey taxon richness to begin to plateau at 51 taxa. With our present sample size, we achieved 98.1% and 76.9% sample coverage for the otter and mink respectively (Fig. 5bii). To achieve 95% sample coverage for the mink, we would need an additional 37 mink scats (54 total). Despite the disparities in sample size, it is unlikely that the mink would consume more prey taxa than the otter (Fig. 5biii).

**Figure 5.**
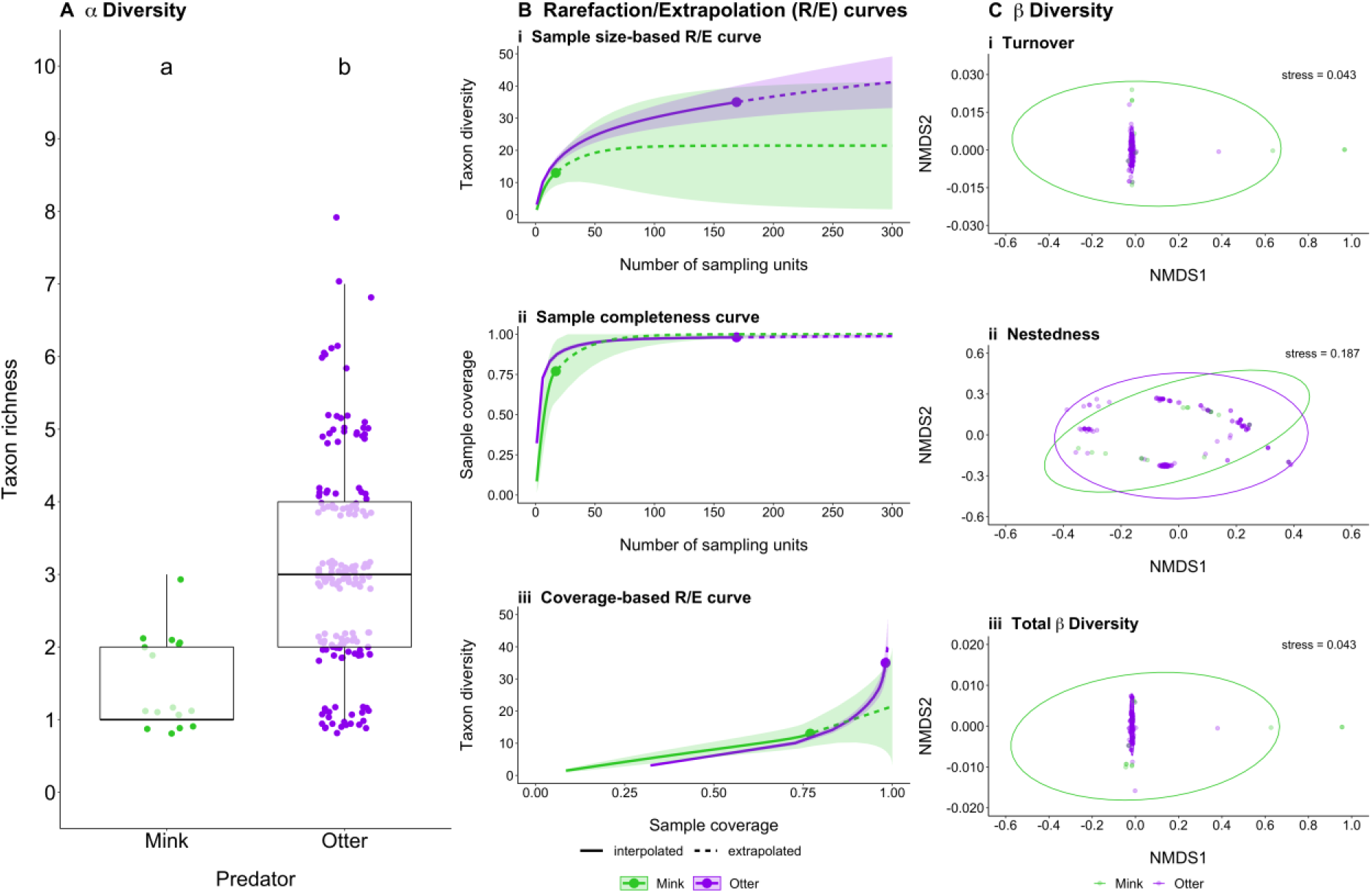
Summaries of alpha and beta diversity comparisons made between otter (purple points/ellipses) and mink (green points/ellipses) faecal samples: **A** boxplot showing the number of prey taxa detected in mink and otter samples, **B** rarefaction/extrapolation (R/E) curves produced for otter spraints and mink scats using iNEXT (Hsieh et al. 2016), and **C** Non-metric Multidimensional Scaling (NMDS) plots of prey communities from otter and mink faecal samples for each beta diversity component. Letters denote significance, where different letters indicate a statistically significant difference in taxon richness. Boxes show 25th, 50th, and 75th percentiles, and whiskers show 5th and 95th percentiles.

Beta diversity of both otter and mink faecal samples was largely driven by turnover (otter: 99.51%; mink: 98.90%) as opposed to nestedness-resultant (otter: 0.49%; mink: 1.10%). MVDISP was different between predators for turnover and total beta diversity, where mink scats had significantly higher dispersion than otter spraints, but not nestedness-resultant (Table 1). Predator had a weak positive influence on the turnover (Fig. 5ci) and total beta diversity (Fig. 5ciii) of prey communities, but not nestedness-resultant (Fig. 5cii; Table 1). Therefore, prey items consumed by the otter were fundamentally different taxa to prey items consumed by the mink, resulting in dissimilar prey community composition.

**Table 1.** Summary of analyses statistically comparing homogeneity of multivariate dispersions between prey communities in otter and mink faecal samples (ANOVA), and variation in prey community composition of otter and mink faecal samples (PERMANOVA).

|  | Homogeneity of multivariate dispersions (ANOVA) |  |  |  | Community similarity (PERMANOVA) |  |  |  |
| --- | --- | --- | --- | --- | --- | --- | --- | --- |
| | Mean distance to centroid ± SE | df | F | P | df | F | R<sup>2</sup> | P |
| *Turnover* |  | 1 | 7.316 | **0.008** | 1 | 5.587 | 0.030 | **0.001** |
| Otter | 0.516 ± 0.031 | | | | | | | |
| Mink | 0.636 ± 0.003 | | | | | | | |
| *Nestedness-resultant* |  | 1 | 0.018 | 0.895 | 1 | -3.097 | -0.017 | 0.915 |
| Otter | 0.107 ± 0.014 | | | | | | | |
| Mink | 0.103 ± 0.006 | | | | | | | |
| *Total beta diversity* |  | 1 | 6.401 | **0.012** | 1 | 4.274 | 0.023 | **0.001** |
| Otter | 0.574 ± 0.014 | | | | | | | |
| Mink | 0.651 ± 0.001 | | | | | | | |

### Geographic variation in otter diet

Of 171 otter spraints, 25 came from Malham Tarn, 38 came from the River Glaven, and 125 came from the River Hull. Two samples (1 each from Malham Tarn and the River Glaven) were removed from the dataset for alpha and beta diversity analyses as they did not contain any prey taxa. Site influenced alpha diversity of otter spraints (χ^2^_2_ = 21.876, *p* < 0.001), where otter spraints from Malham Tarn (*Z* = −3.029, adjusted *p* [Benjamini-Hochberg] = 0.004) and the River Glaven (*Z* = −4.116, adjusted *p* [Benjamini-Hochberg] < 0.001) exhibited lower taxon richness than spraints from the River Hull. Taxon richness in otter spraints from Malham Tarn and the River Glaven did not significantly differ (*Z* = 0.439, adjusted *p* [Benjamini-Hochberg] = 0.661) (Fig. 6a).

**Figure 6.**
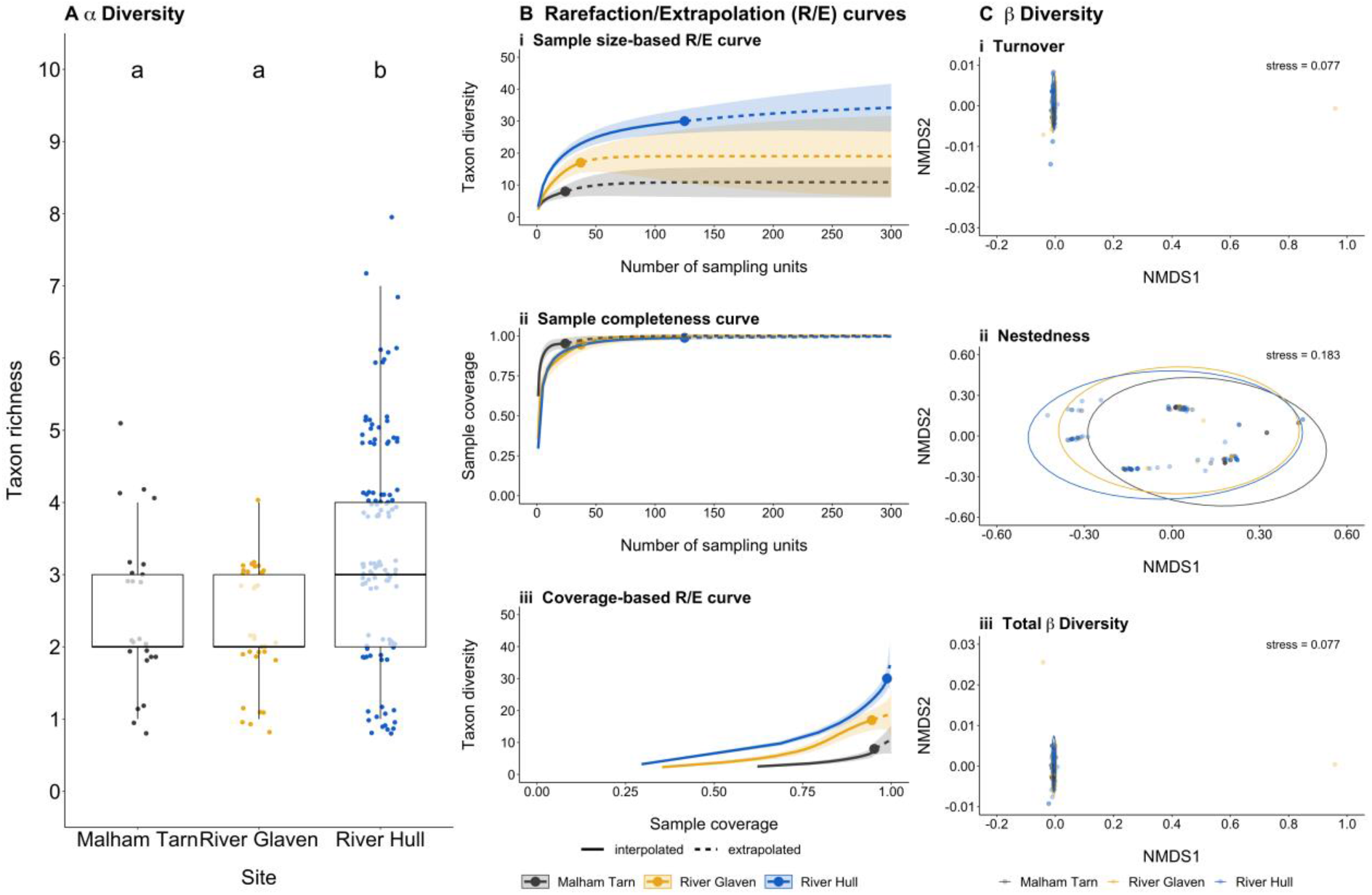
Summaries of alpha and beta diversity comparisons made between otter samples collected from Malham Tarn (grey points/ellipses), River Glaven (yellow points/ellipses), and River Hull (blue points/ellipses): **A** boxplot showing the number of prey taxa detected in samples from each site, **B** rarefaction/extrapolation (R/E) curves produced for otter spraints from Malham Tarn, the River Glaven, and the River Hull using iNEXT (Hsieh et al. 2016), and **C** Non-metric Multidimensional Scaling (NMDS) plots of prey communities in samples from each site for each beta diversity component. Letters denote significance, where different letters indicate a statistically significant difference in taxon richness. Boxes show 25th, 50th, and 75th percentiles, and whiskers show 5th and 95th percentiles.

Rarefaction and extrapolation curves indicated that lower prey taxon richness of Malham Tarn and River Glaven otter spraints was not due to disparities in sample size between sites. Prey taxon richness began to plateau at 10 and 19 taxa with 54 and 107 otter spraints from Malham Tarn and the River Glaven respectively. In contrast, prey taxon richness did not plateau for the River Hull even with 300 otter spraints, at which 38 taxa would be detected (Fig. 6bi). Over 1100 otter spraints from the River Hull would be required for prey taxon richness to begin to plateau at 44 taxa. With our present sample size, we achieved 95.2%, 94.5%, and 98.3% sample coverage for Malham Tarn, the River Glaven, and the River Hull respectively (Fig. 6bii). To achieve 95% sample coverage for the River Glaven, we would need an additional two otter spraints (39 total). Despite the disparities in sample size, it is unlikely that the otter would consume more prey taxa at Malham Tarn or the River Glaven than the River Hull (Fig. 6biii).

Beta diversity of otter samples from all sites was largely driven by turnover (Malham Tarn: 86.91%; River Glaven: 98.41%; River Hull: 99.24%) as opposed to nestedness-resultant (Malham Tarn: 13.09%; River Glaven: 1.59%; River Hull: 0.76%). MVDISP was different between sites for turnover, nestedness-resultant, and total beta diversity, where samples from the River Glaven and River Hull had greater dispersion than samples from Malham Tarn (Table 2). Site had a moderate positive influence on turnover (Fig. 6ci) and weak positive influence on total beta diversity (Fig. 6ciii) of prey communities, but not nestedness-resultant (Fig. 6cii; Table 2). Therefore, prey taxa consumed by otters at a given site were replaced by different prey taxa at other sites.

**Table 2.** Summary of analyses statistically comparing homogeneity of multivariate dispersions between prey communities in otter samples from different sites (ANOVA), and variation in prey community composition of otter samples from different sites (PERMANOVA).

|  | Homogeneity of multivariate dispersions (ANOVA) |  |  |  | Community similarity (PERMANOVA) |  |  |  |
| --- | --- | --- | --- | --- | --- | --- | --- | --- |
| | Mean distance to centroid ± SE | df | F | P | df | F | R<sup>2</sup> | P |
| *Turnover* |  | 2 | 22.620 | **<0.001** | 2 | 10.668 | 0.115 | **0.001** |
| Malham Tarn | 0.220 ± 0.042 | | | | | | | |
| River Glaven | 0.516 ± 0.031 | | | | | | | |
| River Hull | 0.491 ± 0.035 | | | | | | | |
| *Nestedness-resultant* |  | 2 | 11.263 | **<0.001** | 2 | -13.730 | -0.201 | 1.000 |
| Malham Tarn | 0.234 ± 0.028 | | | | | | | |
| River Glaven | 0.079 ± 0.012 | | | | | | | |
| River Hull | 0.117 ± 0.015 | | | | | | | |
| *Total beta diversity* |  | 2 | 23.358 | **<0.001** | 2 | 7.819 | 0.087 | **0.001** |
| Malham Tarn | 0.343 ± 0.052 | | | | | | | |
| River Glaven | 0.564 ± 0.018 | | | | | | | |
| River Hull | 0.560 ± 0.015 | | | | | | | |

## Discussion

We have demonstrated that DNA metabarcoding of otter and mink faeces using vertebrate-specific primers is suitable for dietary assessment, and could be applied to other vertebrate carnivores. We identified a wide range of fish, amphibians, birds, and mammals, all of which were plausible prey items of the otter and mink due to previous species records from each study site. Incorporation of this molecular tool into future dietary assessments for the native otter and invasive mink will enhance our understanding of niche separation between these mustelids.

### Predator assignment

In our study, nearly 10% of scats were misidentified visually and corrected based on predator reads from DNA metabarcoding. Thirteen mink, four fox, and one polecat sample(s) were misidentified as otter spraints. Although collector experience likely influenced this error rate, collectors had received training and most had substantial experience of scat collection for otter diet studies. Similarly, Harrington et al. (2010) found that 75 scats identified as mink by experienced field surveyors actually belonged to pine marten (*Martes martes*), fox, otter, polecat, or stoat (*Mustela erminea*) using DNA barcoding. Scat misidentification can lead to inclusion of prey species consumed by non-focal predators and omission of prey species consumed by the focal predator(s) in dietary assessments, which could have detrimental implications for species conservation and/or management (Martínez-Gutiérrez et al. 2015; Akrim et al. 2018). Therefore, DNA barcoding (Davison et al. 2002; Harrington et al. 2010; Shehzad et al. 2012a, 2012b; Akrim et al. 2018) or DNA metabarcoding (Berry et al. 2017; Forin-Wiart et al. 2018) should be used to identify scats where possible.

Presence of predator DNA is double-edged and can also complicate DNA metabarcoding. Scats from mammalian carnivores can include intact DNA from hairs ingested during grooming (Carss and Parkinson 1996; Shehzad et al. 2012a; Reid et al. 2013) and from intestinal mucosa cells of the defecating predator (Oehm et al. 2011). This can lead to faecal samples being swamped by predator DNA and masking of degraded prey DNA, resulting in reduced detection probability (Shehzad et al. 2012b; Piñol et al. 2015; Robeson et al. 2018; Forin-Wiart et al. 2018; Traugott et al. 2020). This issue can sometimes be alleviated by adding consumer-specific blocking primers (Shehzad et al. 2012a, 2012b; De Barba et al. 2014; Robeson et al. 2018), but potential drawbacks include coblocking of closely related prey taxa, an increased number of sequencing artefacts, and alteration of compositional dietary profiles (Shehzad et al. 2012b; Piñol et al. 2014, 2015; McInnes et al. 2016; Robeson et al. 2018). In our study, otter and mink DNA was present in faecal samples at moderate frequencies (31% and 48% of reads respectively), but did not swamp prey DNA pools acquired for these predators. Higher frequencies of predator DNA were observed in the few fox and polecat samples, but samples still contained a sufficient number of prey reads for reliable identification. Balanced prey and predator DNA in faecal samples is a prerequisite for high detection probability of prey species as well as reliable predator identification, and raises the possibility of using faecal DNA for genotyping individual predators (Bayerl et al. 2017).

### Otter diet

Our finding that otter diet mainly consisted of fish (81.1%), followed by amphibians (12.7%), birds (5.9%) and mammals (0.5%) is consistent with the results of morphological analyses that visually identified prey remains in spraints or stomachs (Jędrzejewska et al. 2001; Clavero et al. 2003; Britton et al. 2006; Reid et al. 2013; Krawczyk et al. 2016; Lanszki et al. 2016). For example, in comparable habitats of the Pannonian biogeographical region, Lanszki et al. (2016) found similar relative occurrence frequencies of fish (82.9%), amphibians (5.1%), reptiles (0.6%), birds (6.7%), mammals (1.0%), crayfish (1.4%), and other invertebrates (2.3%) in otter spraints from rivers, and fish (81.6%), amphibians (7.7%), reptiles (0.8%), birds (2.6%), mammals (1.3%), crayfish (0.4%) and other invertebrates (5.8%) in otter spraints from ponds using morphological analysis. Overall, our results indicate that there was a significant difference in prey community composition of otter spraints at species-level across sites, suggesting that otter diet is highly situational and determined by local variation in prey availability. This is consistent with the wide variety of dietary profiles for the otter reported by previous morphological studies (Ruiz-Olmo et al. 2001; Britton et al. 2006, 2017; Remonti et al. 2010; Reid et al. 2013; Krawczyk et al. 2016; Lanszki et al. 2016). Our results are also in agreement with faecal DNA metabarcoding studies of otter diet. Both Buglione et al. (2020) and Martínez-Abraín et al. (2020) found fish were the primary food resource for otters, followed by amphibians. Specifically, Cyprinidae, Gobidae, Salmonidae, and Percidae were the predominant prey taxa.

Otter diet and fish assemblages in the River Glaven catchment have been extensively studied (Zambrano et al. 2006; Sayer et al. 2011; Almeida et al. 2012, 2013; Champkin et al. 2017). Non-fish species found using morphological spraint analysis included common frog, common toad, grass snake, common moorhen, Eurasian coot, little grebe (*Tachybaptus ruficollis*), mallard (*Anas platyrhynchos*), and water vole (Almeida et al. 2012, 2013). We found that DNA metabarcoding detected all of these species from at least one study site, except for grass snake and little grebe. Several fishes were previously detected by morphological spraint analysis or fish surveys but not by DNA metabarcoding, including stone loach, gudgeon (*Gobio gobio*), ninespine stickleback, ruffe (*Gymnocephalus cernua*), *Lampetra* spp., European flounder (*Platichthys flesus*), rudd (*Scardinius erythrophthalmus*), common bream, goldfish (*Carassius auratus*) and common carp (*Cyprinus carpio*). The common carp and ruffe were initially detected by DNA metabarcoding in agreement with previous morphological studies (Zambrano et al. 2006; Sayer et al. 2011; Almeida et al. 2012; Sayer et al. 2020), but removed by our false positive sequence threshold. Other fishes, while not detected in the River Glaven spraints, were nonetheless detected in spraints from the River Hull or Malham Tarn. The common bream may not have been detected by DNA metabarcoding as this species was last recorded in 1999 by fish surveys at low abundance in one lake (Zambrano et al. 2006). Nondetections of common species in the River Glaven, such as stone loach and brook lamprey (*Lampetra planeri*), may be due to technical bias that can occur throughout the DNA metabarcoding workflow (see *Considerations for molecular scatology*).

Range expansion of the otter into Malham Tarn occurred recently in 2009, and only two individuals have established themselves at the site thus far. Non-fish species found using morphological spraint analysis included common frog, common toad, mallard, tufted duck (*Aythya fuligula*), gull (Laridae spp.), pheasant (*Phasianus colchicus*), and rook (*Corvus frugilegus*) (Alderton et al. 2015). Using DNA metabarcoding, we detected common frog, *Anas* spp., and Laridae spp. in Malham Tarn spraints, and common toad and *Aythya* spp. in River Hull spraints. Fishes detected using morphological spraint analysis or fish surveys included European bullhead, brown trout (*Salmo trutta*), stone loach, perch, and three-spined stickleback. Only brown trout was not detected by DNA metabarcoding at this study site. Large brown trout tend to be open-water feeders in Malham Tarn, whereas juvenile trout reside in the inflow and outflow streams (Eldridge 2016). Absence of brown trout in spraints may reflect a low preference for feeding in open water areas due to the high energy expenditure required to hunt in these habitats in this relatively large lake (Lanszki et al. 2001). In contrast, the European bullhead and stone loach are associated with shoreline cobble-boulder habitats at Malham Tarn, as are small perch (Eldridge 2016). Therefore, habitat associations may explain detection and nondetection of fishes in otter spraints (Lanszki et al. 2001; Alderton et al. 2015).

To our knowledge, no information on otter diet in the River Hull catchment has been published, although research is ongoing (Hänfling et al. unpublished data). Otter diet was most diverse at this site compared to the River Glaven and Malham Tarn, reflecting the higher fish diversity present in this river system. Previous fish surveys of the River Hull using electrofishing or eDNA metabarcoding recorded the same species identified by DNA metabarcoding of otter spraints, except common dace (*Leuciscus leuciscus*), common barbel (*Barbus barbus*), common carp, European chub (*Squalius cephalus*), and tench (*Tinca tinca*). Common carp, common barbel, and European chub were all detected in otter spraints prior to false positive threshold application, but common dace and tench went undetected.

Notwithstanding nondetections at each site, DNA metabarcoding identified species at higher taxonomic resolution than morphological analysis can provide or which morphological identification may miss entirely. Sequences were assigned to common frog and common toad with DNA metabarcoding, whereas amphibian remains are rarely identified to species-level with morphological spraint analysis (Smiroldo et al. 2019). Bird and small mammal remains are typically unidentifiable, or at least challenging to identify, with morphological analysis (Britton et al. 2006; Alderton et al. 2015), yet DNA metabarcoding recorded water vole, common waterfowl (*Anas* spp., *Aythya* spp., Eurasian coot, common moorhen), waders (*Tringa* spp.), gulls (Laridae spp.), and cormorant (*Phalacrocorax carbo*) as well as a number of terrestrial birds, including starling, red-legged partridge (*Alectoris rufa*), stock dove (*Columba oenas*), Eurasian jay (*Garrulus glandarius*), and pheasant. Species-level identification based on morphology is often achievable for smaller fishes (e.g. stickleback species, European bullhead) as otters consume the entire fish resulting in presence of bones in spraints. However, otters only consume selected pieces of flesh and internal organs from larger fishes (e.g. cyprinids, salmonids) and frequently abandon the remainder as an unfinished meal (Almeida et al. 2013). Low occurrence of hard prey components from larger fish in otter spraints may prevent morphological identification, especially of closely related cyprinids (e.g. common carp, goldfish, crucian carp, and their hybrids) which have similar scales. This does not pose an issue for DNA metabarcoding so detection may be improved with molecular scatology.

Despite the regional differences in otter diet, some common dietary patterns emerged. The otter has been reported to selectively predate slow-moving and smaller prey (Chanin 1981; Martínez-Abraín et al. 2019, 2020), with diet reflecting both species and size composition of fish communities occupying their territory. Consistent with previous morphological studies of the River Glaven and Malham Tarn (Almeida et al. 2012; Alderton et al. 2015), we found that otters primarily consumed slow-moving, small species, with less frequent predation on larger species. The European bullhead was the most commonly consumed species at all three study sites. This small benthic species tends to utilise camouflage over escape movements, and it is clear that this strategy may not be effective for avoiding capture by the otter. Additionally, it is possible that the otter has developed unique capture behaviour with regards to European bullhead. Malham Tarn observational work indicated that otters exhibited vigorous rolling and thrashing behaviours in shallow rocky water, presumably to reveal European bullhead presence when hidden amongst cobble-boulder structures (Alderton et al., 2015). Other small, littoral, and benthic species with similar characteristics, such as three-spined stickleback, ninespine stickleback, and stone loach, were also among the most frequently consumed species. Capture of these species might require very little energy expenditure by the otter, even relative to their size, whereas larger, faster fish provide more energy but require more energy to catch and a longer handling time (Remonti et al. 2010; Martínez-Abraín et al. 2019). Therefore, smaller fishes that can be consumed whole are often preferred, although habitat conditions and fish abundance also play a role (Ruiz-Olmo et al. 2001; Britton et al. 2006, 2017; Remonti et al. 2010; Krawczyk et al. 2016; Lanszki et al. 2016; Martínez-Abraín et al. 2019). European bullhead and stickleback species are common at all three of our study sites (Sayer et al. 2011, 2020; Almeida et al. 2012, 2013; Alderton et al. 2015; Champkin et al. 2017; Harwood et al. 2019; Hänfling et al. unpublished data), thus their frequent occurrence in spraints may simply reflect their high abundance in the environment.

Some medium-sized species were also consumed frequently where they were common, such as the European perch in the River Hull catchment and Malham Tarn, and the crucian carp in the River Glaven catchment, a frequent species in farmland ponds (Sayer et al. 2011, 2020). Conversely, other medium-sized or large species which are abundant at our study sites, such as brown trout, common dace, roach and European eel, seemed to be underrepresented in spraints. The fish size categories used here are based on average adult sizes and therefore may not fully explain underrepresentation of these species. Most of these species (apart from European eel) are fast-swimming, open water species, even as juveniles. As such, their capture might require more energy than that of benthic and littoral species. Molecular data cannot reveal the size of individual fish consumed, but morphological spraint analysis has repeatedly shown that small-sized individuals are preferred. For example, a study in South West England showed that European eels of 180 to 270 mm and cyprinids and salmonids of 40 to 130 mm were preferred over larger specimens (up to 440 mm) (Britton et al. 2006). Yet, otters preferred fish weighing between 500-1000 g in a fish pond and streams in the Lake Balaton catchment in Hungary (Lanszki et al. 2001). More detailed quantitative data on fish abundance in the environment are required to distinguish prey selection from density-dependent predation. Indeed, small and benthic species are often underreported in conventional fish surveys, but recent eDNA metabarcoding studies have shown that these species might be much more abundant than previously thought (Hänfling et al. 2016; Li et al. 2019; Griffiths et al. 2020).

Amphibians are an important secondary food resource for otters, comprising up to 43% (average 12%) of otter diet in a meta-analysis of 64 morphological studies conducted across Europe (Smiroldo et al. 2019). Seasonal peaks in otter predation of amphibians tend to coincide with amphibian reproduction in spring and reduced fish availability in winter (Sidorovich 2000; Lanszki et al. 2001; Britton et al. 2006; Prigioni et al. 2006; Reid et al. 2013; Almeida et al. 2013; Alderton et al. 2015; Smiroldo et al. 2019). In our study, occurrence frequency of amphibians in otter diet was on par with previous estimates, particularly common frog and common toad (Jędrzejewska et al. 2001; Clavero et al. 2003; Smiroldo et al. 2019). This was likely due to a high abundance of anurans in ponds next to the River Glaven, River Hull, and Malham Tarn. We also found evidence of predation on great crested newt (*Triturus cristatus*), but detections were negated by our stringent false positive threshold. We did not find any reptiles, but otter predation of grass snake (*Natrix natrix*) in the River Glaven catchment has been recorded by morphological spraint analysis (Almeida et al. 2012). Our study reaffirmed that birds and mammals are of tertiary importance to the otter and these predation events are probably opportunistic (Chanin 1981; Lanszki et al. 2001; Jędrzejewska et al. 2001; Clavero et al. 2003; Prigioni et al. 2006; Krawczyk et al. 2016).

### Mink diet

Published diet assessments for the mink are modest in comparison to the otter. In our study, mink diet was dominated by birds (55.9%) and mammals (39.6%) with only a small component of fish (4.5%). A morphological study in the Biebrza Wetlands of Poland also observed that more mammals (43.7%), fish (32.9%) and birds (21.5%) than amphibians (1.9%) and invertebrates (0.1%) were consumed by the mink in a harsh winter, yet the importance of mammals (68.8%), amphibians (27.2%), birds (1.2%), fish (2.7%) and invertebrates (0.1%) shifted in a mild winter (Skierczyński and Wiśniewska 2010). These results and our own somewhat contrast with other estimates obtained using morphological analyses. Across the Palaearctic region, the mink on average consumed mostly fish (31.9%) and small mammals (25.4%), supplemented by birds (16.2%), amphibians (11.9%), crustaceans (11.0%), and other invertebrates (2.9%) (Jędrzejewska et al. 2001), but consumption varies with location. For example, in woodland streams and rivers of Poland, mink diet was dominated by fish (spring-summer: 40%; autumn-winter: 10%), amphibians (spring-summer: 32%; autumn-winter: 51%), and mammals (spring-summer: 21%; autumn-winter: 36%) (Jędrzejewska et al. 2001). In the Lovat River of Belarus, mink diet was composed of amphibians (ranging from 14-72%, mean 37%) and small mammals (4-80%, mean 27%), supplemented by fish and crayfish (Sidorovich 2000). Despite these overall differences in mink diet, individual prey items found in morphological studies were also identified here, including three-spined stickleback, duck species, Eurasian coot, common moorhen, starling, bank vole, water shrew, brown rat, and European rabbit (Chanin 1981; Jędrzejewska et al. 2001; Bonesi et al. 2004; Bonesi and Macdonald 2004b; Melero et al. 2008; Harrington et al. 2009). Importantly, we also identified water vole in mink scats which is an endangered species in the UK (Mathews and Harrower 2020).

The molecular assay used here does not target invertebrates, but previous morphological studies have shown that these taxa, especially crayfish, can constitute a substantial proportion of otter (average 11.2%) and mink (average 13.9%) diet depending on the biogeographical region studied (Jędrzejewska et al. 2001; Lanszki et al. 2016). For example, the native white-clawed crayfish (*Austropotamobius pallipes*) and invasive signal crayfish (*Pacifastacus leniusculus*) occurred at a frequency of 8.7-25% in otter spraints from the River Glaven catchment (Almeida et al. 2012). The otter and mink may consume more arthropods and molluscs, which are of low energetic value, when fish composition and abundance changes (Clavero et al. 2003; Bonesi et al. 2004). Typical prey species include *Gammarus pulex*, *Asellus aquaticus*, *Dytiscus* spp., white-clawed crayfish, signal crayfish, and the invasive red swamp crayfish (*Procambarus clarkii*) (Carss and Parkinson 1996; Lanszki et al. 2001; Jędrzejewska et al. 2001; Britton et al. 2006; Melero et al. 2008; Almeida et al. 2012, 2013; Reid et al. 2013; Alderton et al. 2015; Martínez-Abraín et al. 2020), but smaller invertebrates could be instances of secondary predation. Future diet assessments for the otter and mink using DNA metabarcoding should also target invertebrates and investigate their role in niche partitioning between these mustelids.

### Niche partitioning between the otter and mink

Our network analysis indicated that the otter used more available resources than the mink and mink diet was less diverse. This is consistent with many other morphological studies which conclude that the otter is a generalist (Prigioni et al. 2006; Remonti et al. 2010) or an opportunist whose diet varies with prey availability and latitude (Clavero et al. 2003; Almeida et al. 2012, 2013; Reid et al. 2013; Alderton et al. 2015), although it has also been called a specialist with respect to diet being limited to aquatic prey such as fish and amphibians (Sidorovich 2000; Bonesi et al. 2004; Bonesi and Macdonald 2004b; Melero et al. 2008; Skierczyński and Wiśniewska 2010; Krawczyk et al. 2016). Conversely, the mink has been observed to utilise both aquatic and terrestrial resources (Sidorovich 2000; Jędrzejewska et al. 2001; Bonesi et al. 2004; Bonesi and Macdonald 2004b; McDonald et al. 2007; Brzeziński et al. 2008; Melero et al. 2008; Skierczyński and Wiśniewska 2010). Results from previous morphological studies (Harrington et al. 2009) and presented here suggest that the mink specialises on terrestrial prey when coexisting with the otter.

With the caveat of a small sample size, we found low niche overlap (0.267) between the otter and mink in our study, which may be indicative of interspecific competition. Mink have been found to consume less fish and more birds and mammals in areas where otters were present, while the otter predominantly consumed fish and amphibians (Chanin 1981; Jędrzejewska et al. 2001; Bonesi et al. 2004; Melero et al. 2008; Harrington et al. 2009). High niche overlap between the mink and otter was found in Poland (Jędrzejewska et al. 2001) and Belarus (Sidorovich 2000), whereas low niche overlap was observed in North East Spain (Melero et al. 2008) using morphological analysis. Niche overlap may vary by geographic region and with predator density, prey composition, season, and environmental conditions (e.g. habitat, weather). In Belarus, higher niche overlap was identified in spring and autumn than summer or winter due to greater availability and consumption of amphibians by both the otter and mink (Sidorovich 2000). In Poland, higher niche overlap was found in spring-summer than autumn-winter (Jędrzejewska et al. 2001), in harsh winter conditions as opposed to mild winter conditions, and in a wetland complex compared to a river catchment (Skierczyński and Wiśniewska 2010). In the UK, Bonesi et al. (2004) found niche overlap between the otter and mink decreased following an increase in otter density and establishment of a resident population, and niche overlap was lower in winter than spring possibly due to resource restrictions. The majority of faecal samples in our study were collected in spring 2015 and autumn 2018, and our results suggest that niche partitioning between the otter and mink may occur year-round.

Importantly, our study was of small geographic extent and analysed few mink scats relative to otter spraints. Across the UK, the native otter is recovering and the subject of ongoing conservation efforts, whereas the invasive mink has declined due to eradication programmes, ongoing control measures, and interspecific aggression from the otter. Therefore, otter spraints are much more abundant and easily sampled than mink scats. Upscaled investigations of otter and mink faeces collected from different freshwater habitats across all seasons are needed to improve understanding of resource use and niche overlap in these mustelids. Despite these limitations, our findings combined with those of previous morphological studies indicate that niche partitioning, through dietary and spatial segregation, between the otter and mink is probable in areas where these mustelids are sympatric and there is an abundance of aquatic and terrestrial resources (Chanin 1981; Bonesi et al. 2004; Bonesi and Macdonald 2004b; Brzeziński et al. 2008; Melero et al. 2008; Harrington et al. 2009). Evidently, the otter and mink can coexist, thus natural biological control of the invasive mink by the native otter will be insufficient on its own to reduce populations of the former. Continued deployment of artificial control methods will be required to eradicate the mink, but biological control can aid these efforts and promote conservation of species impacted by mink activity (Bonesi and Macdonald 2004a, 2004b; Melero et al. 2008; Harrington et al. 2009).

### Considerations for faecal DNA metabarcoding

Bias stemming from choices made throughout the DNA metabarcoding workflow can produce false positive and false negative detections. Scats collected in the field may originate from relatively few individuals, and samples may not be independent (Carss and Parkinson 1996). In the context of our study, male otters have relatively large home ranges (up to 40 km along the length of a river) and return to the same feeding sites (Kruuk 2006). Many of the otter spraints collected from the River Hull catchment may originate from the same territorial male (known from photographs taken by wildlife enthusiasts and trail cameras along the River Hull) that has been present for the last 10 years. Therefore, future DNA metabarcoding studies should include genotyping (Bayerl et al. 2017) and sex-specific markers (Schwarz et al. 2018) to obtain information on identity and sex of predators. This will avoid pseudoreplication (Carss and Parkinson 1996) and provide insights into individual and intersexual variation in diet (Schwarz et al. 2018). Concerning otters, this will also provide information on the communicatory role of sprainting (Kean et al. 2015).

After deposition, scats may be exposed to abiotic and biotic factors that can influence their integrity as well as prey DNA degradation, including temperature (i.e. heat and dehydration), rainfall, UV exposure, coprophagous insects, microbial activity, and decomposition (Carss and Parkinson 1996; Davison et al. 2002; King et al. 2008; Harrington et al. 2010; Oehm et al. 2011; McInnes et al. 2016). Scats may remain in the environment for days or weeks before collection, thus scat freshness is key (Davison et al. 2002; King et al. 2008; De Barba et al. 2014; McInnes et al. 2016). Scats should ideally be collected when an animal is observed defecating, but proxies for freshness include moisture, odour, colour, and consistency (King et al. 2008; McInnes et al. 2016). Scats deposited on vegetation and soil were also found to have lower prey diversity than those deposited on rock or plastic, which may be related to inhibitory compounds present and microbial activity in soil or non-target DNA, e.g. plants, fungi (Oehm et al. 2011; McInnes et al. 2016). In our study, 13 faecal samples (12 otter and one mink according to field identification) failed to produce enough reads for predator assignment and dietary analyses, and another four (two otter, two mink) did not contain any prey taxa. This may be related to freshness or substrate, or these samples may have been deposited by individuals that were fasting due to territorial defence, prey availability, dispersal, pregnancy, rearing young, or limited mobility (McInnes et al. 2016). Future investigations should assess the influence of scat freshness, substrate, and fasting in the otter and mink on prey detection.

Back in the laboratory, DNA extraction may influence prey detection probabilities, including sample coverage, the protocol used (e.g. commercial vs. modular, designed for faeces vs. other substrates) and its efficiency (King et al. 2008; Harrington et al. 2010; Oehm et al. 2011). Prey DNA can be non-uniform in predator faecal samples, thus it may be necessary to subsample or homogenise faeces for DNA extraction (Gosselin et al. 2016) to prevent failed samples. Mustelid scats also contain a number of volatile organic compounds that can be problematic for DNA extraction and PCR (Sellers et al. 2018; Traugott et al. 2020). Both the Qiagen^®^ DNeasy PowerSoil Kit and Mu-DNA soil protocols used here were demonstrated to produce high purity DNA yields from otter spraints suitable for PCR amplification (Sellers et al. 2018). However, we cannot rule out the possibility of DNA degradation or co-extraction of humic substances, phenolic compounds, and proteins in the 13 failed samples. Quality and quantity of prey DNA may be further enhanced by performing extraction replicates for each sample and passing the lysate for each through one spin column or sequencing each independently (King et al. 2008). Extraction, PCR, and sequencing replication also allows occupancy modelling to identify potential false positives arising from secondary predation or contamination and to estimate species detection probabilities (Ficetola et al. 2015).

Secondary predation has been documented in morphological studies of otter spraints and stomachs, where smaller fish consumed by directly predated larger fish inflate prey diversity and bolster the relative importance of small fish as a resource (Carss and Parkinson 1996; Britton et al. 2006), but may be more pronounced in DNA metabarcoding studies due to the greater sensitivity of PCR amplification (Sheppard et al. 2005; King et al. 2008; Pompanon et al. 2012). Secondary predation is challenging to identify in predators that feed on resources at multiple trophic levels, and can affect the inferences made from dietary assessments (Sheppard et al. 2005; Traugott et al. 2020). High sensitivity of DNA metabarcoding also facilitates amplification of contaminants present at minimal concentrations, originating from the environment (e.g. water swallowed with prey, substrate collected with faeces) or the laboratory (King et al. 2008; Pompanon et al. 2012; De Barba et al. 2014; Nielsen et al. 2018; Traugott et al. 2020). Despite physical separation of pre-PCR and post-PCR processes, and common preventative measures for contamination (cleaning workspaces and equipment with 10% bleach solution, filter tips, UV irradiation of plastics and reagents) (King et al. 2008; Pompanon et al. 2012; Traugott et al. 2020), we observed faecal samples were contaminated with our positive control DNA. Error during PCR and sequencing, such as primer mismatch (Piñol et al. 2018) and “tag jumps” (Schnell et al. 2015), can give rise to false positives, cross-contamination between samples, or laboratory contamination (Pompanon et al. 2012). We employed a stringent false positive sequence threshold, which removed false positives from secondary predation or contamination, but also removed potential prey species for the otter and mink that have been reported in previous morphological and metabarcoding studies, e.g. great crested newt (Smiroldo et al. 2019), goldfish (Martínez-Abraín et al. 2020), common carp (Britton et al. 2006; Almeida et al. 2012), and common barbel (Ruiz-Olmo et al. 2001; Britton et al. 2017). This highlights the importance of minimising contamination for lower sequence thresholds and enhanced detection of prey taxa occurring at lower frequencies.

## Conclusions

We have demonstrated the potential of faecal DNA metabarcoding for investigation of diet and niche separation in mustelids as well as predator identification. Despite associated biological and technical challenges, DNA metabarcoding can enhance dietary insights and trophic networks to enable more effective conservation and management of predators and the resources on which they depend. Upscaled, year-round studies on the native otter and invasive mink that screen an equal number of faecal samples for each predator across broader spatial scales, including different freshwater habitats and environmental gradients (e.g. water quality, land-use), will further advance our understanding of resource use and niche overlap in these mustelids. Combining faecal DNA metabarcoding with eDNA metabarcoding of the associated fish fauna will provide further opportunities for more detailed study of prey selection and dietary preferences.

## Author contributions

B.H and T.B conceived and designed the study. R.H and C.S assisted students with faecal sample collection and provided sample metadata. H.V.W performed DNA extractions and R.D constructed libraries for sequencing. L.R.H completed bioinformatic processing of samples, and analysed the data. L.R.H wrote the manuscript, which all authors contributed critically to drafts of and gave final approval for publication.

## Data accessibility

Raw sequence reads have been archived on the NCBI Sequence Read Archive Study: SRP270831; BioProject: PRJNA644190; BioSamples: SAMN15452877-SAMN15453005 [Library 1] and SAMN15455442-SAMN15455596 [Library 2]; SRA accessions: SRR12168859-SRR12168984 [Library 1] and SRR12176017-SRR12176170 [Library 2]). Jupyter notebooks, R scripts and corresponding data have been deposited in a dedicated GitHub repository, which has been permanently archived (https://doi.org/10.5281/zenodo.4252552).

## Acknowledgements

We would like to thank several undergraduate students from the University of Hull for collecting faecal samples from Malham Tarn and the River Hull catchment: Stefan Rooke, Zoe Latham, Nadine Grey, and Alicia Tredell. We are very grateful to the brilliant Terry Linford, Derek Sayer and Peter Bedell for collecting faecal samples from the River Glaven catchment. We also thank Yorkshire Water for contributing to the funding of this study.

## Notes

https://doi.org/10.5281/zenodo.4114361

### Competing Interest Statement

The authors have declared no competing interest.

### Summary of Updates

This version of the manuscript has been updated for resubmission to a journal.

## References

Akrim F, Mahmood T, Max T, Nadeem MS, Qasim S, Andleeb S (2018) Assessment of bias in morphological identification of carnivore scats confirmed with molecular scatology in north-eastern Himalayan region of Pakistan. PeerJ 6: e5262. https://doi.org/10.7717/peerj.5262

Alderton E, Wicker C, Sayer C, Bradley P (2015) The diet of the Malham Tarn otters: understanding the impacts of a native predator. Field Studies. http://fsj.field-studies-council.org/

Almeida D, Copp GH, Masson L, Miranda R, Murai M, Sayer CD (2012) Changes in the diet of a recovering Eurasian otter population between the 1970s and 2010. Aquatic Conservation: Marine and Freshwater Ecosystems 22: 26–35. https://doi.org/10.1002/aqc.1241

Almeida D, Rodolfo N, Sayer CD, Copp GH (2013) Seasonal use of ponds as foraging habitat by Eurasian otter with description of an alternative handling technique for common toad predation. Folia Zoologica 62: 214–221. https://doi.org/10.25225/fozo.v62.i3.a7.2013

Baselga A, Orme CDL (2012) betapart : an R package for the study of beta diversity: Betapart package. Methods in Ecology and Evolution 3: 808–812. https://doi.org/10.1111/j.2041-210X.2012.00224.x

Bayerl H, Kraus RHS, Nowak C, Foerster DW, Fickel J, Kuehn R (2017) Fast and cost-effective single nucleotide polymorphism (SNP) detection in the absence of a reference genome using semideep next-generation Random Amplicon Sequencing (RAMseq). Molecular Ecology Resources 18: 107–117. https://doi.org/10.1111/1755-0998.12717

Berry TE, Osterrieder SK, Murray DC, Coghlan ML, Richardson AJ, Grealy AK, Stat M, Bejder L, Bunce M (2017) DNA metabarcoding for diet analysis and biodiversity: A case study using the endangered Australian sea lion (*Neophoca cinerea*). Ecology and Evolution 7: 5435–5453. https://doi.org/10.1002/ece3.3123

Bonesi L, Chanin P, Macdonald DW (2004) Competition between Eurasian otter *Lutra lutra* and American mink *Mustela vison* probed by niche shift. Oikos 106: 19–26. https://doi.org/10.1111/j.0030-1299.2004.12763.x

Bonesi L, Macdonald DW (2004a) Impact of released Eurasian otters on a population of American mink: a test using an experimental approach. Oikos 106: 9–18. https://doi.org/10.1111/j.0030-1299.2004.13138.x

Bonesi L, Macdonald DW (2004b) Differential habitat use promotes sustainable coexistence between the specialist otter and the generalist mink. Oikos 106: 509–519. https://doi.org/10.1111/j.0030-1299.2004.13034.x

Bonesi L, Palazon S (2007) The American mink in Europe: Status, impacts, and control. Biological Conservation 134: 470–483. https://doi.org/10.1016/j.biocon.2006.09.006

Britton JR, Pegg J, Shepherd JS (2006) Revealing the prey items of the otter *Lutra lutra* in South West England using stomach contents analysis. Folia Zoologica 55: 167–174.

Britton RJ, Berry M, Sewell S, Lees C, Reading P (2017) Importance of small fishes and invasive crayfish in otter *Lutra lutra* diet in an English chalk stream. Knowledge and management of aquatic ecosystems 418: 13. https://doi.org/10.1051/kmae/2017004

Bronner IF, Quail MA, Turner DJ, Swerdlow H (2009) Improved Protocols for Illumina Sequencing. Current Protocols in Human Genetics 18: 18.2.1–18.2.42. https://doi.org/10.1002/0471142905.hg1802s62

Brzeziński M, Święcicka-Mazan A, Romanowski J (2008) Do Otters and Mink Compete for Access to Foraging Sites? A Winter Case Study in the Mazurian Lakeland, Poland. Annales Zoologici Fennici 45: 317–322. https://doi.org/10.5735/086.045.0412

Buglione M, Petrelli S, Troiano C, Notomista T, Rivieccio E, Fulgione D (2020) The diet of otters (*Lutra lutra*) on the Agri river system, one of the most important presence sites in Italy: a molecular approach. PeerJ 8: e9606. http://doi.org/10.7717/peerj.9606

Carss DN, Parkinson SG (1996) Errors associated with otter *Lutra lutra* faecal analysis. I. Assessing general diet from spraints. Journal of Zoology 238: 301–317. https://doi.org/10.1111/j.1469-7998.1996.tb05396.x

Champkin J, Copp GH, Sayer CD, Clilverd H, Walker AM (2017) Responses of fishes and lampreys to the re-creation of meanders in a small English chalk stream. River Research and Applications 34: 34–43. https://doi.org/10.1002/rra.3216

Chanin P (1981) The diet of the otter and its relations with the feral mink in two areas of southwest England. Acta Theriologica 26: 83–95. https://doi.org/10.4098/AT.arch.81-5

Clavero M, Prenda J, Delibes M (2003) Trophic diversity of the otter (*Lutra lutra* L.) in temperate and Mediterranean freshwater habitats. Journal of Biogeography 30: 761–769. https://doi.org/10.1046/j.1365-2699.2003.00865.x

Davison A, Birks JDS, Brookes RC, Braithwaite TC, Messenger JE (2002) On the origin of faeces: morphological versus molecular methods for surveying rare carnivores from their scats. Journal of Zoology 257: 141–143. https://doi.org/10.1017/S0952836902000730

Deagle BE, Thomas AC, McInnes JC, Clarke LJ, Vesterinen EJ, Clare EL, Kartzinel, TR, Eveson, JP (2018) Counting with DNA in metabarcoding studies: how should we convert sequence reads to dietary data? Molecular Ecology 28: 391–406. https://doi.org/10.1111/mec.14734

De Barba M, Miquel C, Boyer F, Mercier C, Rioux D, Coissac E, Taberlet P (2014) DNA metabarcoding multiplexing and validation of data accuracy for diet assessment: application to omnivorous diet. Molecular Ecology Resources 14: 306–323. https://doi.org/10.1111/1755-0998.12188

Dormann CF, Frueund J, Bluethgen N, Gruber B (2009) Indices, graphs and null models: analyzing bipartite ecological networks. The Open Ecology Journal 2: 7–24. https://doi.org/10.2174/1874213000902010007

Eldridge T (2016) Determining diet interactions of brown trout (*Salmo trutta*) and Eurasian perch (*Perca fluviatilis*) in an upland marl lake using stable isotope analysis. MSc Thesis, University College London.

Ficetola GF, Pansu J, Bonin A, Coissac E, Giguet-Covex C, De Barba M, Gielly L, Lopes CM, Boyer F, Pompanon F, Rayé G, Taberlet P (2015) Replication levels, false presences and the estimation of the presence/absence from eDNA metabarcoding data. Molecular Ecology Resources 15: 543–556. https://doi.org/10.1111/1755-0998.12338

Forin-Wiart M-A, Poulle M-L, Piry S, Cosson J-F, Larose C, Galan M (2018) Evaluating metabarcoding to analyse diet composition of species foraging in anthropogenic landscapes using Ion Torrent and Illumina sequencing. Scientific Reports 8: 17091. https://doi.org/10.1038/s41598-018-34430-7

Gosselin EN, Lonsinger RC, Waits LP (2017) Comparing morphological and molecular diet analyses and fecal DNA sampling protocols for a terrestrial carnivore: Noninvasive Diet Analysis Methodologies. Wildlife Society Bulletin 41: 362–369. https://doi.org/10.1002/wsb.749

Griffiths, NP, Bolland J, Wright R, Murphy LA, Donnelly RK, Watson HV, Hänfling B (2020) Environmental DNA metabarcoding provides enhanced detection of the European eel *Anguilla anguilla* and fish community structure in pumped river catchments. Under review at Journal of Fish Biology.

Hänfling B, Lawson Handley L, Read DS, Hahn C, Li J, Nichols P, Blackman, RC, Oliver A, Winfield IJ (2016) Environmental DNA metabarcoding of lake fish communities reflects long-term data from established survey methods. Molecular Ecology 25: 3101–3119. https://doi.org/10.1111/mec.13660

Hӓnfling B, Lawson Handley L, Harper LR, Benucci M, Sellers GS, Di Muri C, Griffiths NP, Jaques R, James J (2020) Development of an eDNA-based lake fish classification tool – Collection of data from English Lakes. Final Report, February 2020. Written for the Environment Agency, UK (unpublished).

Harper LR, Handley LL, Carpenter AI., Ghazali M, Di Muri C, Macgregor CJ, Logan TW, Law A, Breithaupt T, Read DS, McDevitt AD, Hänfling B (2019a) Environmental DNA (eDNA) metabarcoding of pond water as a tool to survey conservation and management priority mammals. Biological Conservation 238: 108225. https://doi.org/10.1016/j.biocon.2019.108225

Harper LR, Lawson Handley L, Hahn C, Boonham N, Rees HC, Lewis E, Adams IP, Brotherton P, Phillips S, Hänfling B (2019b) Generating and testing ecological hypotheses at the pondscape with environmental DNA metabarcoding: A case study on a threatened amphibian. Environmental DNA 2: 184–199. https://doi.org/10.1002/edn3.57

Harrington LA, Harrington AL, Yamaguchi N, Thom MD, Ferreras P, Windham TR, Macdonald DW (2009) The impact of native competitors on an alien invasive: temporal niche shifts to avoid interspecific aggression? Ecology 90: 1207–1216. https://doi.org/10.1890/08-0302.1

Harrington LA, Harrington AL, Hughes J, Stirling D, Macdonald DW (2010) The accuracy of scat identification in distribution surveys: American mink, *Neovison vison*, in the northern highlands of Scotland. European Journal of Wildlife Research 56: 377–384. https://doi.org/10.1007/s10344-009-0328-6

Harwood A, Berridge R, Perrow M, Piper A, Sayer, CD (2019) Understanding European eel (*Anguilla anguilla*) ecology within a Norfolk river catchment to inform eel management. ENG2083, Draft Final Report, August 2019.

Hsieh TC, Ma KH, Chao A (2016) iNEXT: an R package for rarefaction and extrapolation of species diversity (Hill numbers). Methods in Ecology and Evolution 7: 1451–1456. https://doi.org/10.1111/2041-210X.12613

Jędrzejewska B, Sidorovich VE, Pikulik MM, Jędrzejewska W (2001) Feeding habits of the otter and the American mink in Białowieża Primeval Forest (Poland) compared to other Eurasian populations. Ecography 24: 165–180. https://www.jstor.org/stable/3683692

Kean EF, Chadwick EA, Mueller CT (2015) Scent signals individual identity and country of origin in otters. Mammalian Biology 80: 99–105. https://doi.org/10.1016/j.mambio.2014.12.004

King RA, Read DS, Traugott M, Symondson WOC (2008) Molecular analysis of predation: a review of best practice for DNA-based approaches. Molecular Ecology 17: 947–963. https://doi.org/10.1111/j.1365-294x.2007.03613.x

Krawczyk AJ, Bogdziewicz M, Majkowska K, Glazaczow A (2016) Diet composition of the Eurasian otter *Lutra lutra* in different freshwater habitats of temperate Europe: a review and meta-analysis. Mammal Review 46: 106–113. https://doi.org/10.1111/mam.12054

Kruuk H (2006) Otters: ecology, behaviour, and conservation. Oxford University Press, Oxford.

Lanszki J, Körmendi S, Hancz C, Martin TG (2001) Examination of some factors affecting selection of fish prey by otters (*Lutra lutra*) living by eutrophic fish ponds. Journal of Zoology 255: 97–103. https://doi.org/10.1017/S0952836901001145

Lanszki J, Lehoczky I, Kotze A, Somers MJ (2016) Diet of otters (*Lutra lutra*) in various habitat types in the Pannonian biogeographical region compared to other regions of Europe. PeerJ 4: e2266. https://doi.org/10.7717/peerj.2266

Li J, Hatton-Ellis TW, Lawson Handley L-J, Kimbell HS, Benucci M, Peirson G, Hänfling B (2019) Ground-truthing of a fish-based environmental DNA metabarcoding method for assessing the quality of lakes. Journal of Applied Ecology 56: 1232–1244. https://doi.org/10.1111/1365-2664.13352

Martínez-Abraín A, Santidrián Tomillo P, Veiga J (2019) Otter diet changes in a reservoir during a severe autumn drought. Journal of Mammalogy 101: 211–215. https://doi.org/10.1093/jmammal/gyz185

Martínez-Abraín A, Marí-Mena N, Vizcaíno A, Vierna J, Veloy C, Amboage M, Guitián-Caamaño A, Key C, Vila M (2020) Determinants of Eurasian otter (*Lutra lutra*) diet in a seasonally changing reservoir. Hydrobiologia 847: 1803–1816. https://doi.org/10.1007/s10750-020-04208-y

Martínez-Gutiérrez PG, Palomares F, Fernández N (2015) Predator identification methods in diet studies: uncertain assignment produces biased results? Ecography 38: 922–929. https://doi.org/10.1111/ecog.01040

Mathews F, Harrower C (2020) Regional Red List of British Mammals. The Mammal Society. https://www.mammal.org.uk/science-research/red-list/

McDonald RA, O’Hara K, Morrish DJ (2007) Decline of invasive alien mink (*Mustela vison*) is concurrent with recovery of native otters (*Lutra lutra*). Diversity and Distributions 13: 92–98. https://doi.org/10.1111/j.1366-9516.2006.00303.x

McInnes JC, Alderman R, Deagle BE, Lea M-A, Raymond B, Jarman SN (2016) Optimised scat collection protocols for dietary DNA metabarcoding in vertebrates. Methods in Ecology and Evolution 8: 192–202. https://doi.org/10.1111/2041-210X.12677

McInnes JC, Jarman SN, Lea M-A, Raymond B, Deagle BE, Phillips RA, Catry P, Stanworth A, Weimerskirch H, Kusch A, Gras M, Cherel Y, Maschette D, Alderman R (2017) DNA Metabarcoding as a Marine Conservation and Management Tool: A Circumpolar Examination of Fishery Discards in the Diet of Threatened Albatrosses. Frontiers in Marine Science 4: 277. https://doi.org/10.3389/fmars.2017.00277

Melero Y, Palazón S, Bonesi L, Gosàlbez J (2008) Feeding habits of three sympatric mammals in NE Spain: the American mink, the spotted genet, and the Eurasian otter. Acta Theriologica 53: 263–273. https://doi.org/10.1007/BF03193123

Nakagawa S, Cuthill IC (2007) Effect size, confidence interval and statistical significance: a practical guide for biologists. Biological Reviews 82: 591–605 https://doi.org/10.1111/j.1469-185X.2007.00027.x

Nielsen JM, Clare EL, Hayden B, Brett MT, Kratina P (2018) Diet tracing in ecology: Method comparison and selection. Methods in Ecology and Evolution 9: 278–291. https://doi.org/10.1111/2041-210X.12869

Oehm J, Juen A, Nagiller K, Neuhauser S, Traugott M (2011) Molecular scatology: how to improve prey DNA detection success in avian faeces? Molecular Ecology Resources 11: 620–628. https://doi.org/10.1111/j.1755-0998.2011.03001.x

Ogle DH, Wheeler P, Dinno A (2020) FSA: Fisheries Stock Analysis. R package version 0.8.30. https://github.com/droglenc/FSA

Piñol J, Mir G, Gomez-Polo P, Agustí N (2015) Universal and blocking primer mismatches limit the use of high-throughput DNA sequencing for the quantitative metabarcoding of arthropods. Molecular Ecology Resources 15: 819–830. https://doi.org/10.1111/1755-0998.12355

Piñol J, San Andrés V, Clare EL, Mir G, Symondson WOC (2014) A pragmatic approach to the analysis of diets of generalist predators: the use of next-generation sequencing with no blocking probes. Molecular Ecology Resources 14: 18–26. https://doi.org/10.1111/1755-0998.12156

Piñol J, Senar MA, Symondson WOC (2018) The choice of universal primers and the characteristics of the species mixture determines when DNA metabarcoding can be quantitative. Molecular Ecology 28: 407–419. https://doi.org/10.1111/mec.14776

Pompanon F, Deagle BE, Symondson WOC, Brown DS, Jarman SN, Taberlet P (2012) Who is eating what: diet assessment using next generation sequencing. Molecular Ecology 21: 1931–1950. https://doi.org/10.1111/j.1365-294X.2011.05403.x

Prigioni C. Balestrieri A, Remonti L, Gargaro A, Priore G (2006) Diet of the Eurasian otter (*Lutra lutra*) in relation to freshwater habitats and alien fish species in southern Italy. Ethology Ecology & Evolution 18: 307–320. https://doi.org/10.1080/08927014.2006.9522698

Reid N, Thompson D, Hayden B, Marnell F, Montgomery WI (2013) Review and quantitative meta-analysis of diet suggests the Eurasian otter (*Lutra lutra*) is likely to be a poor bioindicator. Ecological indicators 26: 5–13. https://doi.org/10.1016/j.ecolind.2012.10.017

Remonti L, Prigioni C, Balestrieri A, Sgrosso S, Priore G (2010) Eurasian otter (*Lutra lutra*) prey selection in response to a variation of fish abundance. Italian Journal of Zoology 77: 331–338. https://doi.org/10.1080/11250000903229809

Reynolds JC, Short MJ, Leigh RJ (2004) Development of population control strategies for mink *Mustela vison*, using floating rafts as monitors and trap sites. Biological Conservation 120: 533–543. https://doi.org/10.1016/j.biocon.2004.03.026

Riaz T, Shehzad W, Viari A, Pompanon F, Taberlet P, Coissac E (2011) ecoPrimers: inference of new DNA barcode markers from whole genome sequence analysis. Nucleic Acids Research 39: e145. https://doi.org/10.1093/nar/gkr732

Robeson MS 2nd, Khanipov K, Golovko G, Wisely SM, White MD, Bodenchuck M, Smyser TJ, Fofanov Y, Fierer N, Piaggio, AJ (2018) Assessing the utility of metabarcoding for diet analyses of the omnivorous wild pig (*Sus scrofa*). Ecology and Evolution 8: 185–196. https://doi.org/10.1002/ece3.3638

Ruiz-Olmo J, López-Martín JM, Palazón S (2001) The influence of fish abundance on the otter (*Lutra lutra*) populations in Iberian Mediterranean habitats. Journal of Zoology 254: 325–336. https://doi.org/10.1017/S095283690100083

Sayer CD, Copp GH, Emson D, Godard MJ, Zięba G, Wesley KJ (2011) Towards the conservation of crucian carp *Carassius carassius*: understanding the extent and causes of decline within part of its native English range. Journal of Fish Biology 79: 1608–1624. https://doi.org/10.1111/j.1095-8649.2011.03059.x

Sayer CD, Emson D, Patmore IR, Greaves H, Wiseman G, West WP, Tarkan AS, Davies GD, Payne J, Cooper B, Grapes T, Copp GH (2020) Recovering the crucian carp *Carassius carassius* (L.): approach and early results of an English conservation project. Aquatic Conservation: Marine and Freshwater Ecosystems. https://doi.org/10.1002/aqc.3422

Schnell IB, Bohmann K, Gilbert MTP (2015) Tag jumps illuminated--reducing sequence-to-sample misidentifications in metabarcoding studies. Molecular Ecology Resources 15: 1289–1303. https://doi.org/10.1111/1755-0998.12402

Schwarz D, Spitzer SM, Thomas AC, Kohnert CM, Keates TR, Acevedo-Gutiérrez A (2018) Large-scale molecular diet analysis in a generalist marine mammal reveals male preference for prey of conservation concern. Ecology and Evolution 8: 9889–9905. https://doi.org/10.1002/ece3.4474

Sellers GS, Di Muri C, Gómez A, Hänfling B (2018) Mu-DNA: a modular universal DNA extraction method adaptable for a wide range of sample types. Metabarcoding and Metagenomics 2: e24556. https://doi.org/10.3897/mbmg.2.24556

Shehzad W, McCarthy TM, Pompanon F, Purevjav L, Coissac E, Riaz T, Taberlet P (2012a) Prey preference of snow leopard (*Panthera uncia*) in South Gobi, Mongolia. PLoS ONE 7: e32104. https://doi.org/10.1371/journal.pone.0032104

Shehzad W, Riaz T, Nawaz MA, Miquel C, Poillot C, Shah SA, Pompanon F, Coissac E, Taberlet P (2012b) Carnivore diet analysis based on next-generation sequencing: application to the leopard cat (*Prionailurus bengalensis*) in Pakistan. Molecular Ecology 21: 1951–1965. https://doi.org/10.1111/j.1365-294X.2011.05424.x

Sheppard SK, Bell J, Sunderland KD, Fenlon J, Skervin D, Symondson WOC (2005) Detection of secondary predation by PCR analyses of the gut contents of invertebrate generalist predators. Molecular Ecology 14: 4461–4468. https://doi.org/10.1111/j.1365-294X.2005.02742.x

Sidorovich VE (2000) Seasonal variation in the feeding habits of riparian mustelids in river valleys of NE Belarus. Acta Theriologica 45: 233–242. https://doi.org/10.4098/AT.arch.00-25

Skierczyński M, Wiśniewska A (2010) Trophic niche comparison of American mink and Eurasian otter under different winter conditions. Mammalia 74: 433–437. https://doi.org/10.1515/mamm.2010.043

Smiroldo G, Villa A, Tremolada P, Gariano P, Balestrieri A, Delfino M (2019) Amphibians in Eurasian otter *Lutra lutra* diet: osteological identification unveils hidden prey richness and male-biased predation on anurans. Mammal Review 49: 240–255. https://doi.org/10.1111/mam.12155

Traugott M, Thalinger B, Wallinger C, Sint D (2020) Fish as predators and prey: DNA-based assessment of their role in food webs. Journal of Fish Biology. https://doi.org/10.1111/jfb.14400

Wickham H (2009) ggplot2: Elegant Graphics for Data Analysis. Springer, New York.

Zambrano L, Perrow MR, Sayer CD, Tomlinson ML, Davidson TA (2006) Relationships between fish feeding guild and trophic structure in English lowland shallow lakes subject to anthropogenic influence: implications for lake restoration. Aquatic Ecology 40: 391–405. https://doi.org/10.1007/s10452-006-9037-3

