## Supplementary Material for "Using DNA metabarcoding to investigate diet and niche partitioning in the native European otter (*Lutra lutra*) and invasive American mink (*Neovison vison*)"

#### Contents

Appendix 1: Fish inventory 3
Appendix 2: Methods 4
2.1 DNA metabarcoding 4
2.2 Data analysis 6
Appendix 3: Predator Assignment 8
Appendix 4: Samples from non-focal mammal predators 9
Appendix 5: Non-focal mammal diet 10
Supplementary Tables 11
Table S1 11
Supplementary Figures 12
Figure S1 12
Figure S2 13
Figure S3 14
Figure S4 15
Figure S5 16
Figure S6 17
References 18

#### Appendix 1: Fish inventory

Microsoft Excel spreadsheet (*Appendix1_fish_inventory.xlsx*) detailing the basic inventory of fishes created from available survey data to permit a broad comparison between prey detected in otter spraints by DNA metabarcoding and available prey species. A 'x' inside a yellow box is used to highlight species that were detected by fish surveys but not by faecal DNA metabarcoding. The full website for the Environment Agency database used is as follows: https://data.gov.uk/dataset/f49b8e4b-8673-498e-bead-98e6847831c6/freshwater-fish-counts-for-all-species-all-areas-and-all-years

### Appendix 2: Methods

#### 2.1 DNA metabarcoding

A two-step PCR protocol was performed on faecal samples at the University of Hull. Dedicated rooms were available for pre-PCR and post-PCR processes. Pre-PCR processes were performed in a dedicated eDNA laboratory, with separate rooms for filtration, DNA extraction, and PCR preparation of sensitive environmental samples. PCR reactions were set up in a ultraviolet (UV) and bleach sterilised laminar flow hood. To minimise cross-contamination risk between samples, eight-strip PCR tubes with individually attached lids were used instead of 96-well plates (Port et al. 2016). After PCR reagents excluding template DNA had been added to PCR tubes, the tubes were sealed and transported to a separate laboratory on a different floor for the addition of faecal DNA extracts and the PCR positive control. PCR positive and negative controls were included on each PCR run to screen for sources of potential contamination. The DNA (0.05 ng/µL) used for the PCR positive control was *Maylandia zebra* as this is an exotic cichlid not found in UK freshwater habitats. The negative controls substituted sterile molecular grade water (Fisher Scientific UK Ltd, Loughborough, UK) for template DNA.

During the first PCR, the target region was amplified using published 12S ribosomal RNA (rRNA) primers 12S-V5-F (5'-ACTGGGATTAGATACCCC-3') and 12S-V5-R (5'-TAGAACAGGCTCCTCTAG-3') (Riaz et al. 2011) that were validated *in silico, in vitro,* and *in situ* for all UK vertebrates (Hänfling et al. 2016; Harper et al. 2019a, 2019b). Primers were modified to include indexes, heterogeneity spacers, sequencing primers, and pre-adapters. During the first PCR, three replicates were performed for each sample to combat PCR stochasticity. PCR reactions were performed in 25 µL volumes, consisting of 3 µL of template DNA, 1.5 µL of each 10 µM primer (Integrated DNA Technologies, Belgium), 12.5 µL of Q5® High-Fidelity 2x Master Mix (New England Biolabs® Inc., MA, USA) and 6.5 µL molecular grade water. PCR was performed on an Applied Biosystems® Veriti Thermal Cycler (Life Technologies, CA, USA) with the following thermocycling profile: 98 °C for 5 mins, 35 cycles of 98 °C for 10 s, 58 °C for 20 s and 72 °C for 30 s, followed by a final elongation step at 72 °C for 7 mins. PCR products were stored at 4 °C until replicates for each sample were pooled, and 2 µL of pooled PCR product was added to 0.5 µL of 5x DNA Loading Buffer Blue (Bioline®, London, UK). PCR product was visualised on 2% agarose gels (1.6 g Bioline® Agarose in 80 mL 1x sodium borate) (Brody & Kern, 2004) stained with GelRed® (Cambridge Bioscience, Cambridge, UK), and gels were imaged using Image Lab Software (Bio-Rad Laboratories Ltd, Watford, UK). A PCR product was deemed positive where there was an amplification band on the gel that was of the expected size (200-300 bp). PCR products were stored at -20 °C until they were pooled according to PCR plate to create sub-libraries for purification with Mag-BIND® RxnPure Plus magnetic beads (Omega Bio-tek Inc, GA, USA), following the double size selection protocol established by Bronner et al. (2009). Ratios of 0.9x and 0.15x magnetic beads to 100 µL of each sub-library were used. Eluted DNA (30 µL) was stored at -20 °C until the second PCR could be performed.

The second PCR bound pre-adapters, indexes, and Illumina adapters to the purified sub-libraries. Two replicates were performed for each sub-library in 50 µL volumes, consisting of 6 µL of template DNA, 3 µL of each 10 µM primer (Integrated DNA Technologies, Belgium), 25 µL of Q5® High-Fidelity 2x Master Mix, and 13 µL molecular grade water. PCR was performed on an Applied Biosystems® Veriti Thermal Cycler with the following thermocycling profile: 95 °C for 3 mins, 8 cycles of 98 °C for 20 s and 72 °C for 1 min, followed by a final elongation step at 72 °C for 5 mins. PCR products were stored at 4 °C until duplicates for each sub-library were pooled, and 2 µL of pooled product was added to 0.5 µL of 5x DNA Loading Buffer Blue. PCR products were visualised on 2% agarose gels (1.6 g Bioline® Agarose in 80 mL 1x sodium borate) stained with GelRed®, and gels were imaged using Image Lab Software. Again, PCR products were deemed positive where there was an amplification band on the gel that was of the expected size (300-400 bp). Sub-libraries were stored at 4 °C until purification with Mag-BIND® RxnPure Plus magnetic beads, following the double size selection protocol established by Bronner et al. (2009). Ratios of 0.7x and 0.15x magnetic beads to 50 µL of each sub-library were used. Eluted DNA (30 µL) was stored at 4 °C until normalisation and final purification.

Sub-libraries were quantified on a Qubit™ 3.0 fluorometer using a Qubit™ dsDNA HS Assay Kit (Invitrogen, UK) and pooled proportional to sample size and concentration. Each pooled library was purified using the same ratios, volumes, and protocol as second PCR purification. Based on Qubit™ concentration, the libraries were diluted to 4 nM for quantification by real-time quantitative PCR (qPCR) using the NEBNext® Library Quant Kit for Illumina® (New England Biolabs® Inc., MA, USA) on a StepOnePlus™ Real-Time PCR system (Life Technologies, CA, USA). The libraries were also checked using an Agilent 2200 TapeStation and High Sensitivity D1000 ScreenTape (Agilent Technologies, CA, USA) to verify secondary product had been removed successfully and a fragment of the expected size (330 bp) remained. The libraries were sequenced at 12 pM with 10% PhiX Control v3 on an Illumina MiSeq® using a MiSeq Reagent Kit v3 (600-cycle) (Illumina Inc., CA, USA).

Raw sequence reads were demultiplexed using a custom Python script then processed using metaBEAT (metaBarcoding and Environmental Analysis Tool) v0.97.11 (https://github.com/HullUni-bioinformatics/metaBEAT). Raw reads were quality trimmed from the read ends (minimum per base phred score Q30) and across sliding windows (window size 5bp; minimum average phred score Q30) using Trimmomatic v0.32 (Bolger et al. 2014). Reads were cropped to a maximum length of 110 bp and reads shorter than 90 bp after quality trimming were discarded. The first 18 bp of remaining reads were also removed to ensure no locus primer remained. Sequence pairs were merged into single high quality reads using FLASH v1.2.11 (Magoč and Salzberg 2011), provided there was a minimum overlap of 10 bp and no more than 10% mismatch between pairs. Only forward reads were kept for pairs that could not be merged. A final length filter was applied to ensure sequences reflected the expected fragment size (90-110 bp). Retained sequences were screened for chimeric sequences against a custom reference database for UK vertebrates (Harper et al. 2019b) using the uchime algorithm (Edgar et al. 2011), as implemented in vsearch v1.1 (Rognes et al. 2016). Redundant sequences were removed by clustering at 100% identity ('--cluster_fast' option) in vsearch v1.1 (Rognes et al. 2016). Clusters were considered sequencing error and omitted from further processing if they were represented by less than three sequences. Non-redundant sets of query sequences were then compared against the UK vertebrate reference database (Harper et al. 2019b) using BLAST (Zhang et al. 2000). Putative taxonomic identity was assigned using a lowest common ancestor (LCA) approach based on the top 10% BLAST matches for any query that matched a reference sequence across more than 80% of its length at minimum identity of 98%. Unassigned sequences were subjected to a separate BLAST search against the complete NCBI nucleotide (nt) database at 98% identity to determine the source via LCA as described above. The bioinformatic analysis has been deposited in the GitHub repository for reproducibility (permanently archived at: https://doi.org/10.5281/zenodo.4252552).

#### 2.2 Data analysis

Assignments from different databases were merged, and spurious assignments (i.e. species that do not occur in the study area, invertebrates and bacteria) were removed from the dataset. Spurious assignments included: Atlantic herring (*Clupea harengus*), a marine fish; eastern mudminnow (*Umbra pygmaea*), a freshwater fish that does not occur in the UK; mourning dove (*Zenaida macroura*), a North American bird; and the *Pan* genus, which contains two extant African species. Next, higher taxonomic assignments (i.e. genus, family, order) containing only one UK species were reassigned to that species. The order primates was reassigned to human (*Homo sapiens*). The families Cichlidae, Cottidae, Felidae, and Hominidae were reassigned to zebra mbuna (*Maylandia zebra*), European bullhead (*Cottus gobio*), domestic cat (*Felis catus*), and human respectively. The genera *Cottus*, *Meleagris*, and *Vulpes* were reassigned to European bullhead, turkey (*Meleagris gallopavo*), and red fox (*Vulpes vulpes*) respectively. The species *Canis lupus* and *Sus scrofa* were reassigned to domestic dog (*Canis lupus familiaris*) and domestic pig (*Sus scrofa domesticus*) given the restricted distribution of wild boar (*S. scrofa*) and absence of grey wolf (*C. lupus*) in the UK.

The only confirmed misassignment was the cichlid *Rhamphochromis esox* which was reassigned to zebra mbuna. A potential misassignment was green-winged teal (*Anas carolinensis*) which is a rare migrant that has been infrequently recorded in the UK (British Trust for Ornithology, 2019) but may have been assigned due to potential for hybridisation within the genus *Anas* or identicality to mallard (*Anas platyrhynchos*) and Eurasian teal (*Anas crecca*) across the 12S rRNA fragment. This species was reassigned to the genus *Anas*. Reads from corrected assignments were then merged with unaltered assignments. Contamination was observed in PCR positive and negative controls (Fig. S2). Consequently, we applied a sequence threshold, which was the maximum sequence frequency of cichlid DNA in faecal samples (1.123%), to minimise the risk of false positives in our dataset. Taxa were only classed as present in faeces if their sequence frequency exceeded the set threshold, which narrowed detections considerably (Fig. S3).

Assignments higher than species level were then removed from the dataset with the following exceptions. Taxa assigned to genus-level (*Microtus* spp., *Tringa* spp.), but not to individual species with reference sequence data, were treated as a single taxon.

For genera and families where species could not be unambiguously distinguished using our reference sequence database or several species were missing reference sequences, all assignments were pooled at the genus-level (*Anas* spp., *Aythya* spp.) or family-level (Laridae spp., Percidae spp.) and treated as a single taxon. Human and domestic animals (cow [*Bos taurus*], dog, pig) were regarded as environmental contaminants and also removed for the purposes of downstream analyses. Therefore, all taxonomic assignments in the final dataset were predominantly of species resolution and considered real detections.

#### Appendix 3: Predator assignment

Microsoft Excel spreadsheet (*Appendix3_predator_assignment.xlsx*) used to identify the mammal predator for each faecal sample based on the proportional predator read counts. Mammal predator read counts in each sample were summed, and the proportional read counts for each predator species were calculated from the total predator read counts.

#### Appendix 4: Samples from non-focal mammal predators

Microsoft Excel spreadsheet (*Appendix4_fox_polecat_analysis.xlsx*) used to analyse red fox (*Vulpes vulpes*) and European polecat (*Mustela putorius*) samples. The total percentage of prey (by vertebrate group) sequences relative to predator sequences was evaluated across all samples belonging to each predator.

#### Appendix 5: Non-focal mammal diet

A total of three and one prey taxa were detected in the fox and polecat samples respectively. Fox DNA and polecat DNA was more abundant in faecal samples from these predators at 77.6% and 97.1% respectively, and only other mammals were predated (Supplementary Material: Appendix 4). Using the prey reads, *Microtus* spp. (33.8%), bank vole (*Myodes glareolus*) (1.8%), and European rabbit (*Oryctolagus cuniculus*) (64.4%) comprised fox diet. *Microtus* spp. and European rabbit were each detected in two fox faecal samples and bank vole identified from one fox faecal sample. For polecat, only *Microtus* spp. (100%) was detected in the single faecal sample from this predator.

#### Supplementary Tables

**Table S1.** Sample information, including collection date, coordinates, and site, made available as an excel spreadsheet (*TableS1_sample_metadata.xlsx*).

#### Supplementary Figures

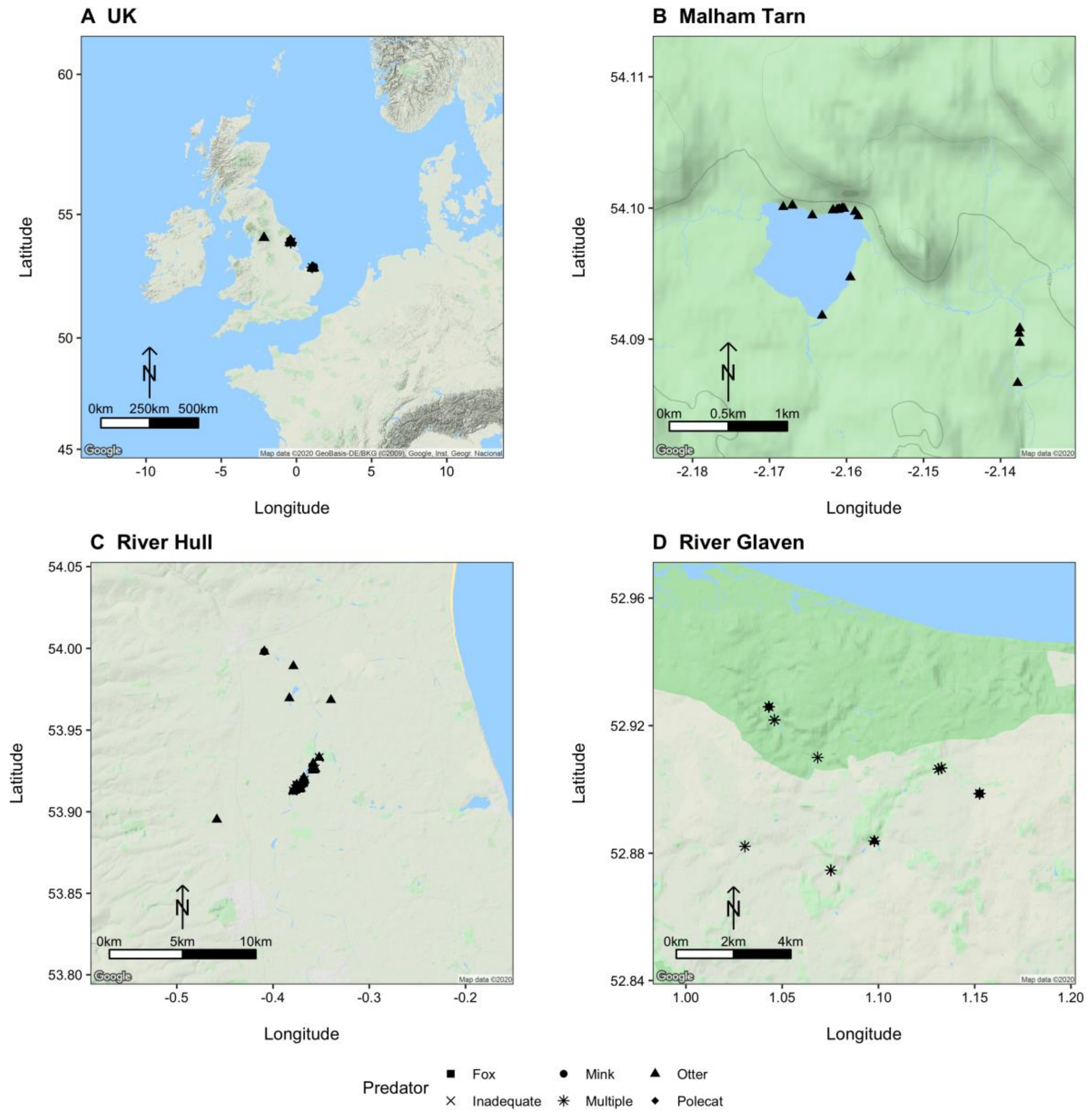


**Figure S1.** Maps showing sampling locations across the UK and at each study site: **A** UK, **B** Malham Tarn, West Yorkshire, **C** River Hull catchment, East Yorkshire, and **D** River Glaven catchment, Norfolk. Mammal predator for each sample based on spraint morphology upon collection is represented by different shapes. 'Inadequate' samples refer to samples that had <100 reads for any mammal predator. 'Multiple' samples refer to samples that contained reads belonging to multiple mammal predators.

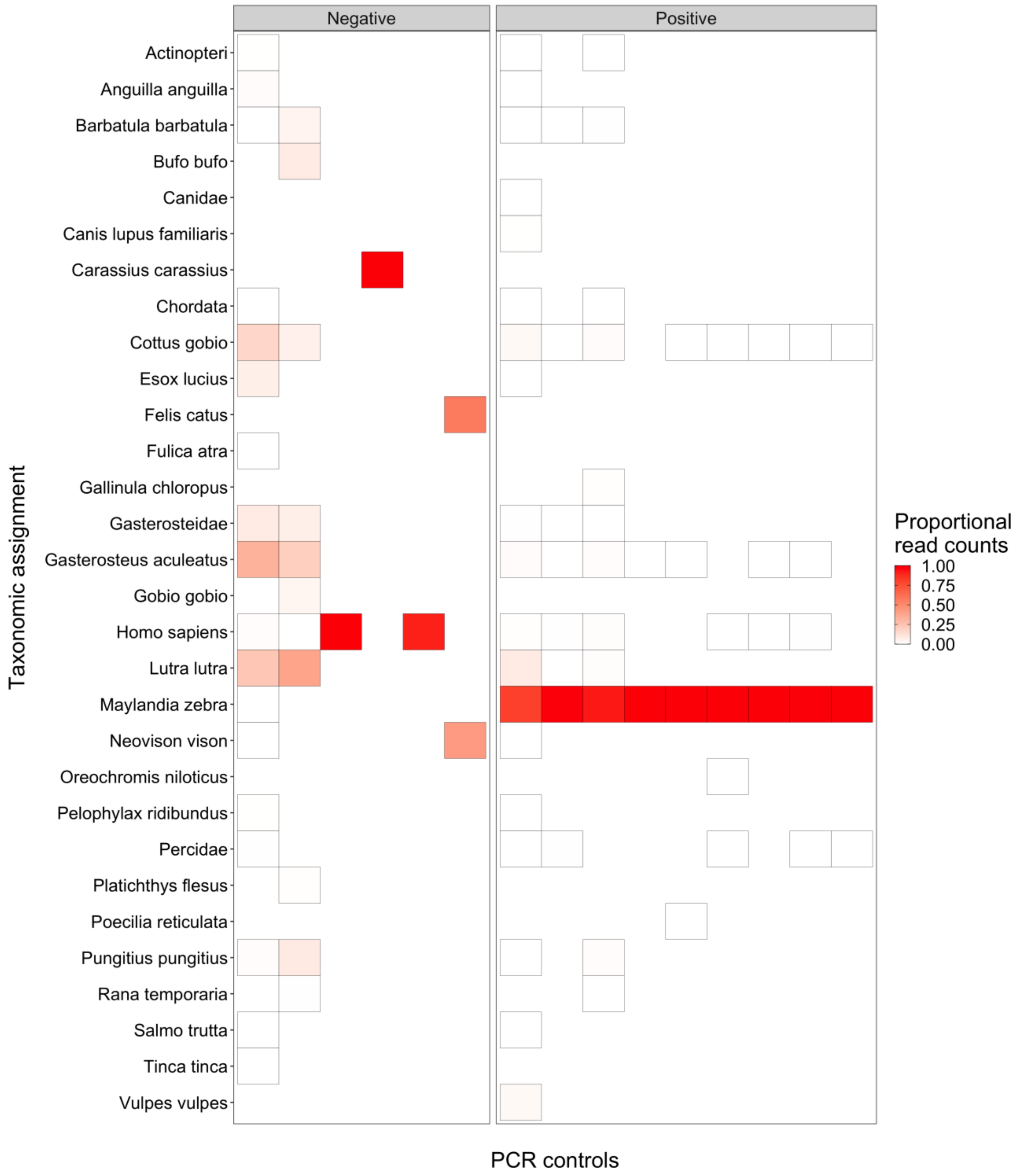


**Figure S2.** Heatmap showing the frequency of contamination in PCR negative controls (molecular grade water) and PCR positive controls (*Maylandia zebra*). Assignments that were not detected in a PCR control are represented by white tiles with no border.

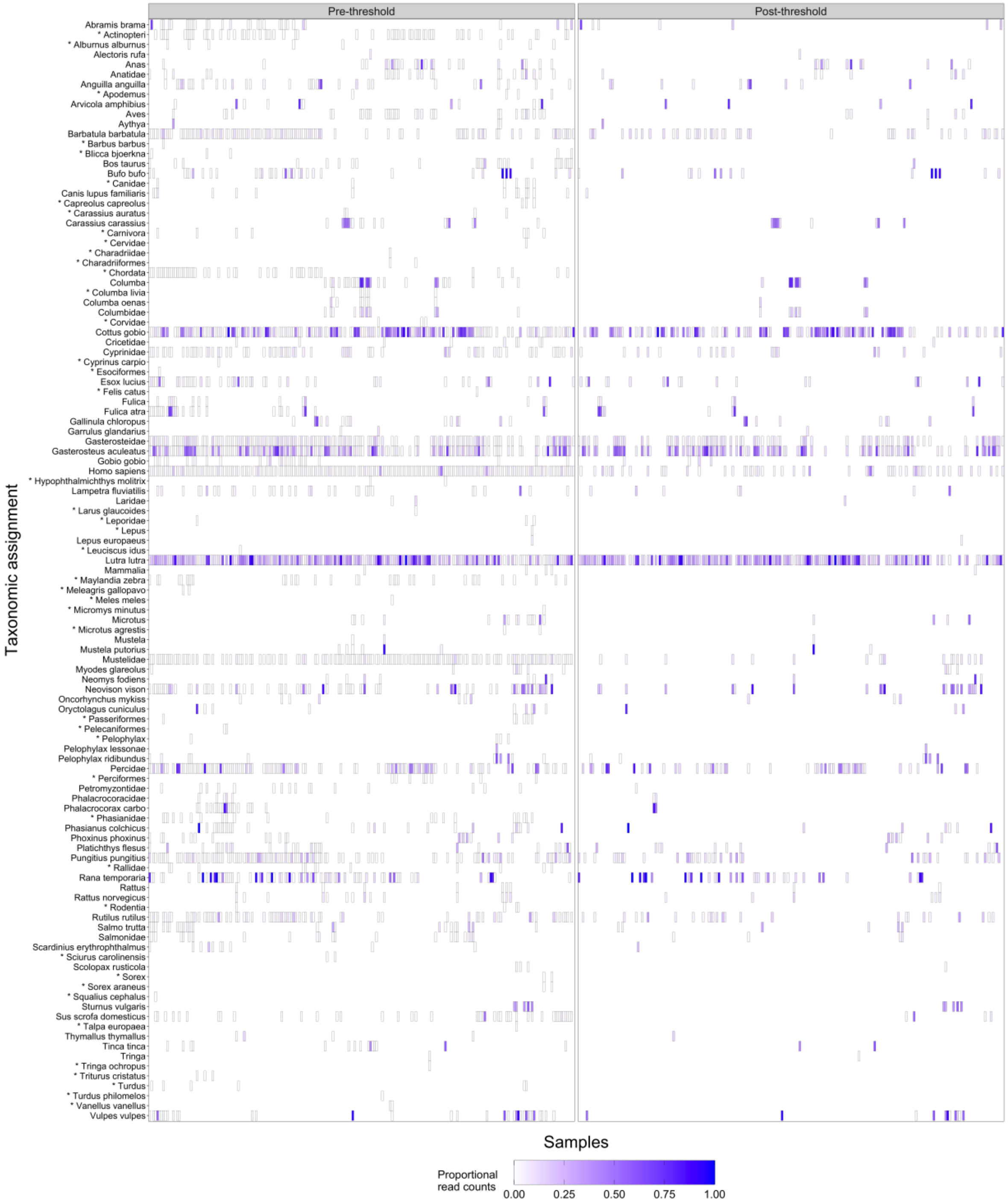


**Figure S3.** Heatmaps showing proportional read counts for taxa found in faecal samples before and after false positive sequence threshold application. Taxa that were removed by the false positive threshold are highlighted with an asterisk. The PCR positive control (*Maylandia zebra*) was not found in any faecal samples after threshold application. Taxa that were not detected in a sample are represented by white tiles with no border.

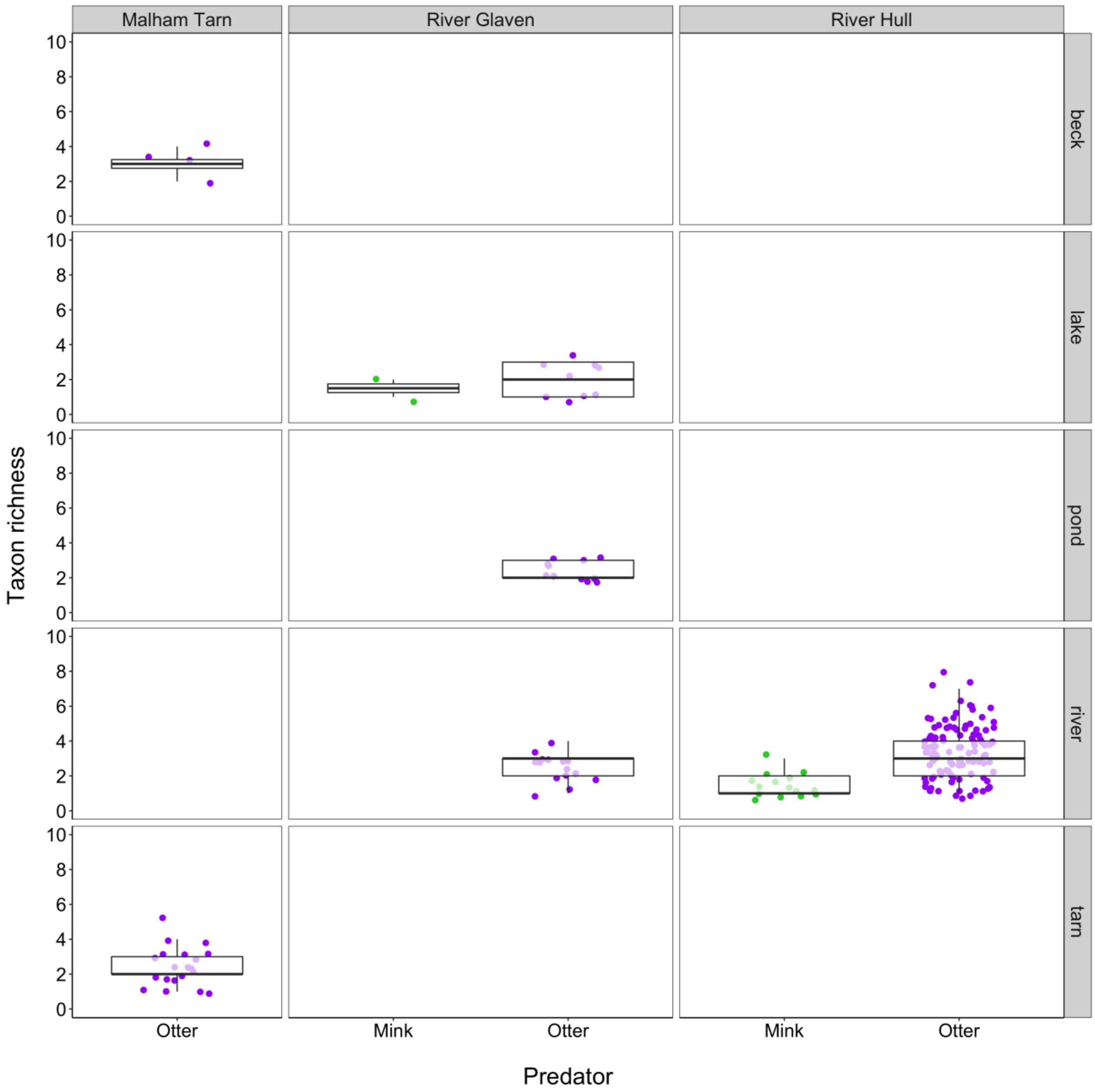


**Figure S4.** Boxplot of prey taxon richness for each predator according to site and waterbody where faecal samples were collected. Boxes show 25th, 50th, and 75th percentiles, and whiskers show 5th and 95th percentiles.

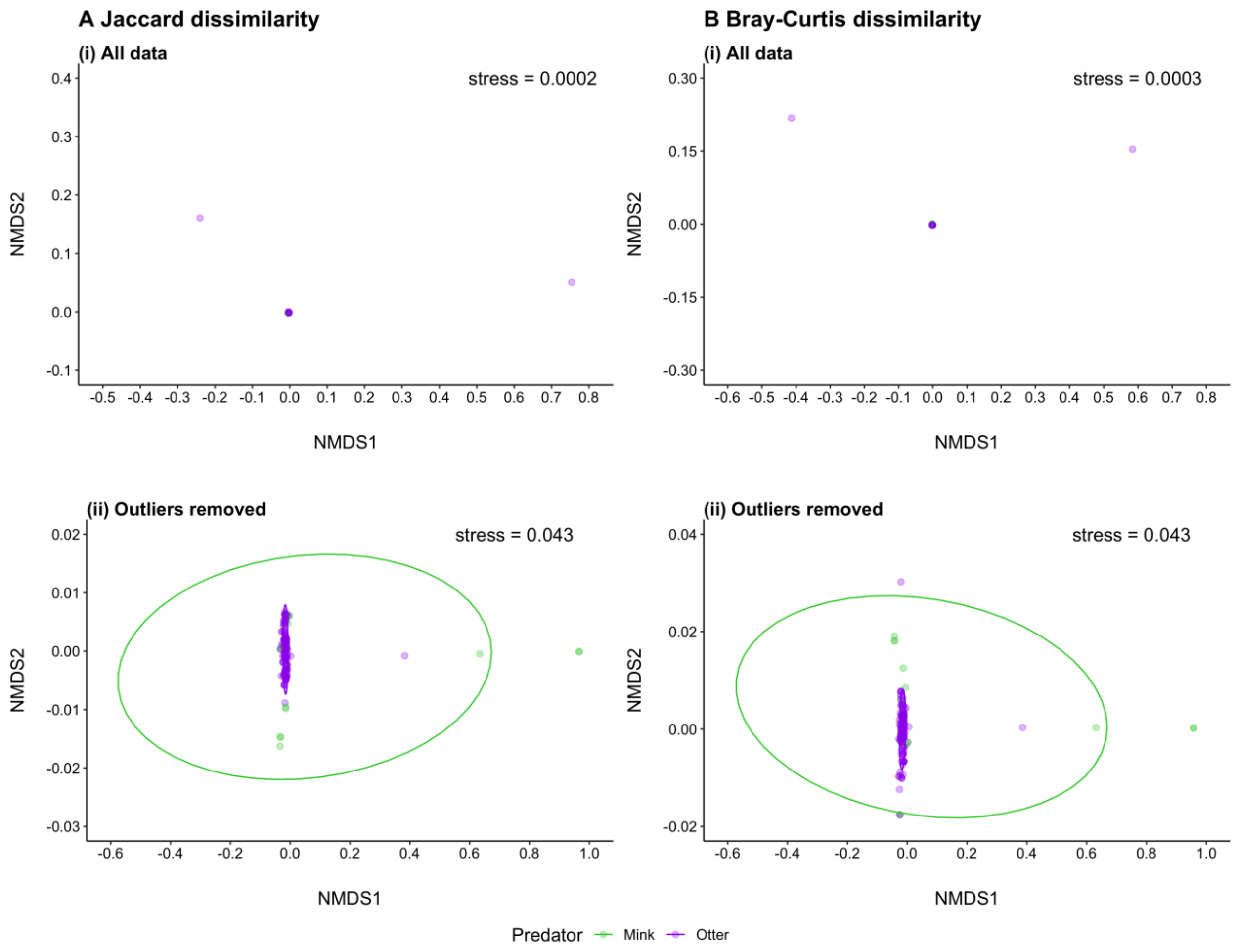


**Figure S5.** Plots summarising three-dimensional Non-metric Multidimensional Scaling (NMDS) of prey communities from otter and mink faecal samples based on: **A** occurrence data (Jaccard dissimilarity), and **B** relative read abundance data (Bray-Curtis dissimilarity). Data were analysed including all faecal samples **(i)** and excluding extreme outliers **(ii)**.

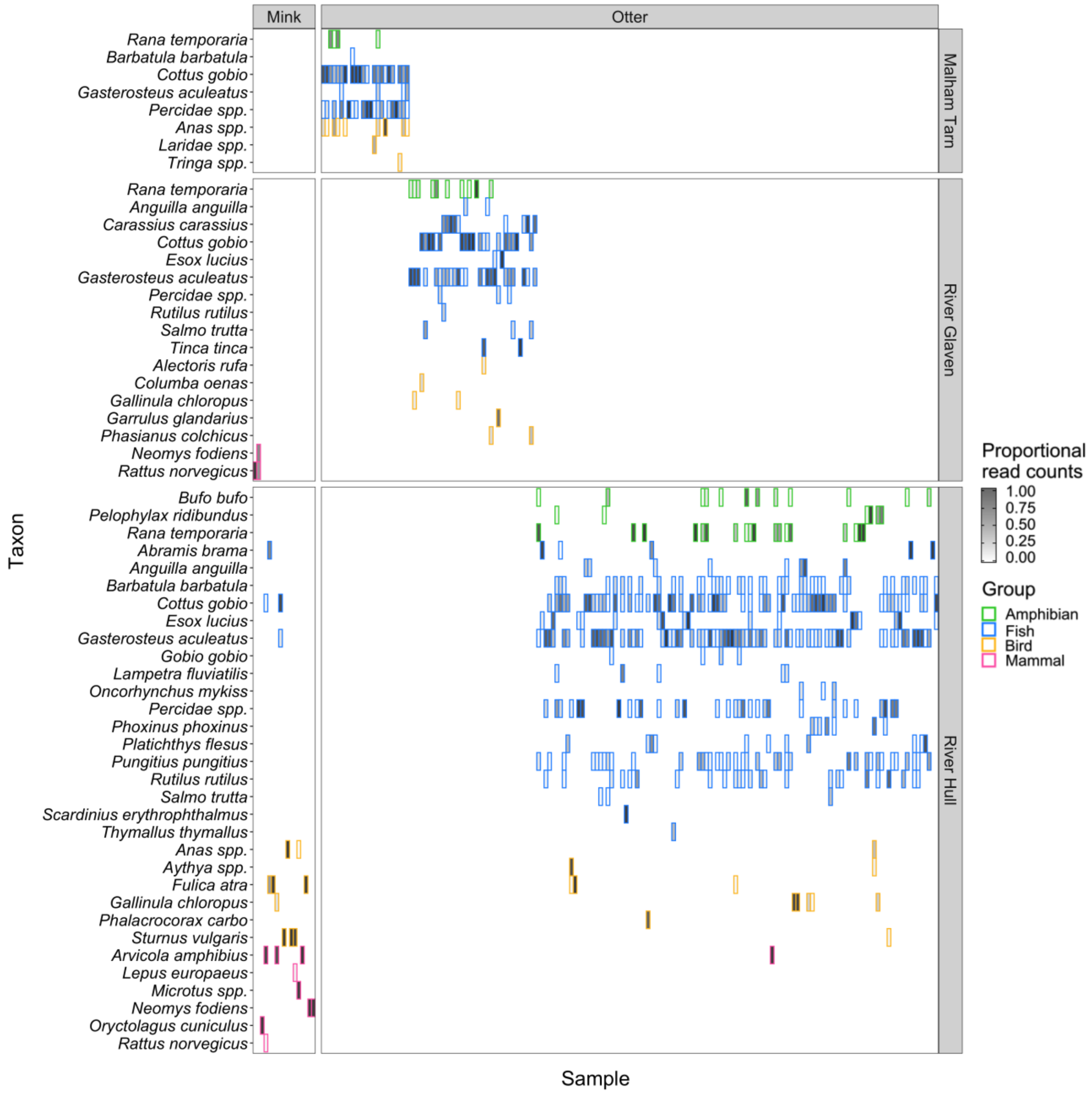


**Figure S6.** Heatmap showing proportional read counts for prey taxa detected in otter and mink faeces. Tile borders are coloured according to vertebrate group. Taxa that were not detected in a faecal sample are represented by white tiles with no border.

#### References


Bolger AM, Lohse M, Usadel B (2014) Trimmomatic: a flexible trimmer for Illumina sequence data. Bioinformatics 30: 2114–2120. https://doi.org/10.1093/bioinformatics/btu170

Brody JR, Kern SE (2004) Sodium boric acid: a Tris-free, cooler conductive medium for DNA electrophoresis. BioTechniques 36: 214–216. https://doi.org/10.2144/04362BM02

Bronner IF, Quail MA, Turner DJ, Swerdlow H (2009) Improved Protocols for Illumina Sequencing. Current Protocols in Human Genetics 18: 18.2.1–18.2.42. https://doi.org/10.1002/0471142905.hg1802s80

Edgar RC, Haas BJ, Clemente JC, Quince C, Knight R (2011) UCHIME improves sensitivity and speed of chimera detection. Bioinformatics 27: 2194–2200. https://doi.org/10.1093/bioinformatics/btr381

Hänfling B, Lawson Handley L, Read DS, Hahn C, Li J, Nichols P, Blackman RC, Oliver A, Winfield, IJ (2016) Environmental DNA metabarcoding of lake fish communities reflects long-term data from established survey methods. Molecular Ecology 25: 3101–3119. https://doi.org/10.1111/mec.13660

Harper LR, Handley LL, Carpenter AI., Ghazali M, Di Muri C, Macgregor CJ, Logan TW, Law A, Breithaupt T, Read DS, McDevitt AD, Hänfling B (2019a) Environmental DNA (eDNA) metabarcoding of pond water as a tool to survey conservation and management priority mammals. Biological Conservation 238: 108225. https://doi.org/10.1016/j.biocon.2019.108225

Harper LR, Lawson Handley L, Hahn C, Boonham N, Rees HC, Lewis E, Adams IP, Brotherton P, Phillips S, Hänfling B (2019b) Generating and testing ecological hypotheses at the pondscape with environmental DNA metabarcoding: A case study on a threatened amphibian. Environmental DNA 2: 184–199. https://doi.org/10.1002/edn3.57

Magoč T, Salzberg SL (2011) FLASH: fast length adjustment of short reads to improve genome assemblies. Bioinformatics 27: 2957–2963. https://doi.org/10.1093/bioinformatics/btr507

Port JA, O'Donnell JL, Romero-Maraccini OC, Leary PR, Litvin SY, Nickols KJ, Yamahara KM, Kelly, RP (2016) Assessing vertebrate biodiversity in a kelp forest ecosystem using environmental DNA. Molecular Ecology 25: 527–541. https://doi.org/10.1111/mec.13481

Riaz T, Shehzad W, Viari A, Pompanon F, Taberlet P, Coissac E (2011) ecoPrimers: inference of new DNA barcode markers from whole genome sequence analysis. Nucleic Acids Research 39: e145. https://doi.org/10.1093/nar/gkr732

Rognes T, Flouri T, Nichols B, Quince C, Mahé F (2016) VSEARCH: a versatile open source tool for metagenomics. PeerJ 4: e2584. https://doi.org/10.7717/peerj.2584

Zhang Z, Schwartz S, Wagner L, Miller W (2000) A greedy algorithm for aligning DNA sequences. Journal of Computational Biology 7: 203–214. https://doi.org/10.1089/10665270050081478
